# The human telomeric nucleosome displays distinct structural and dynamic properties

**DOI:** 10.1101/2019.12.18.881755

**Authors:** Aghil Soman, Chong Wai Liew, Hsiang Ling Teo, Nikolay V. Berezhnoy, Vincent Olieric, Nikolay Korolev, Daniela Rhodes, Lars Nordenskiöld

## Abstract

Telomeres protect the ends of our chromosomes and are key to maintaining genomic integrity during cell division and differentiation. However, our knowledge of telomeric chromatin and nucleosome structure at the molecular level is limited. Here, we aimed to define the structure, dynamics as well as properties in solution of the human telomeric nucleosome. We first determined the 2.2 Å crystal structure of a human telomeric nucleosome core particle (NCP) containing 145 bp DNA, which revealed the same helical path for the DNA as well as symmetric stretching in both halves of the NCP as that of the 145 bp ‘601’ NCP. In solution, the telomeric nucleosome exhibited a less stable and a markedly more dynamic structure compared to NCPs containing DNA positioning sequences. These observations provide molecular insights into how telomeric DNA forms nucleosomes and chromatin and advance our understanding of the unique biological role of telomeres.

## INTRODUCTION

In the nucleus of eukaryotic cells, DNA is packaged into chromatin, comprising a linear array of DNA-histone protein complexes called nucleosomes. Nucleosomes are the structural subunits of the chromosomes and their physical and chemical properties determine multiple aspects of gene translation, genomic stability and accessibility to regulatory protein binding. The central part of the nucleosome is the nucleosome core particle (NCP) consisting of ∼146 bp of DNA wrapped in a 1.75-turn super-helix around one histone octamer (HO), which in turn is composed of two copies of each of the four histone proteins H2A, H2B, H3 and H4 (1,2). The formation of the NCP, and therefore of chromatin, is driven by electrostatic interactions and requires the DNA helix to bend and wrap tightly around the histone octamer. Since the structure and flexibility of DNA is dependent on its sequence, it follows that different DNA sequences will require different energies to wrap around the histone octamer. Consequently, genome-wide analyses showed that nucleosome positioning (and occupancy) is DNA sequence-dependent (3-6). It was earlier shown by *in vitro* analysis of the DNA extracted from chicken erythrocyte NCPs that AT rich TA/AT and AA/TT base steps are favoured in positions where the DNA minor groove faces inwards contacting the HO (3). Additionally, GC rich base steps preferentially locate where the major groove is facing inward (3,7).

Mammalian telomeres, the ends of linear chromosomes, are uniquely made of highly conserved TTAGGG DNA sequence repeats that are hotspots for DNA damage (8,9). Telomeres act as a platform for the recruitment of telomere-specific DNA binding factors that form the protective shelterin complex (10-13). These specialised regions are therefore critical for maintaining genomic integrity by protecting the ends of the eukaryotic chromosomes from incorrect DNA repair and degradation (10-13), and accordingly aberrant regulation of telomeres has been linked with tumour development and pathologies of ageing (14,15). Despite this, we know almost nothing about the structure of telomeric nucleosomes at the molecular level. Previous studies showed that telomeric DNA forms nucleosomes and is chromatinised, but with an unusually short nucleosome repeat length of about 157 bp (16-18). Telomeric DNA lacks positioning information due to the repetitive nature of its hexameric DNA sequence, which is out of phase with the DNA helical repeat that is close to 10 bp (19). Likely, this is the cause of the observed weak affinity for HO binding by telomeric DNA (16,20-22), leading to dynamic telomeric nucleosomes (23,24). What remains unclear is how the apparent conflict between the repetitive nature of the telomeric sequence and the sequence demands of nucleosome formation is resolved, and how/whether the unique sequence properties of telomeres are linked to their key biological functions.

Our knowledge on the nucleosomal structure of chromatin comes primarily from crystal structures of NCPs. Finch et al. obtained the first crystals of NCPs isolated from native chromatin (2), but all subsequent high-resolution structural studies on nucleosomes have used core particles originating from three DNA nucleosome-positioning sequences. The strongest known is the Widom ‘601’ sequence obtained from systematic evolution of ligands by exponential enrichment (SELEX) experiments performed *in vitro* (25,26). These results specifically highlighted the important role of the most flexible of all base steps, the TA step, occurring with a 10 bp periodicity at inward minor groove registers. The only positioning DNA sequences of natural origin (albeit not fully native NCPs) for which NCP crystal structures have been determined are the centromeric α-satellite and the mouse mammary tumor virus (MMT-V) promoter sequence (27-29). Two cryo-EM structures of nucleosome formed by native positioning DNA and the native human histone octamer have been solved recently (30,31). The question of whether natural and biologically important non-positioning sequences like telomeric DNA, forms stable and well-defined nucleosome cores as we know them, is therefore still an open issue.

Here we aimed to define the structure, dynamics and properties in solution of the human telomeric NCP, in order to answer the question of how telomeric DNA forms nucleosomes and how this might relate to the known biological functions of the telomere. We determined the 2.2 Å crystal structure of a telomeric nucleosome core particle (Telo-NCP) containing 145 bp human telomeric DNA comprising 23 TTAGGG repeats, reconstituted with human histones, and asked questions about its stability and dynamics in solution.

## MATERIALS AND METHODS

### DNA and histone octamer preparations

The plasmid containing the monomer of a 145 bp of telomeric DNA (5′-ATC-(TTAGGG)_23_TGAT-3′) flanked by *EcoRV* and *AvaI* sites was obtained from Bio Basic Asia Pacific Pte Ltd. A plasmid containing eight copies of this 145 bp telomeric DNA was cloned into Sure2 *E.coli* (Agilent Technologies Singapore Pte. Ltd) following established protocols (32). The resulting plasmid was grown at 30 °C for 18 hours in TB medium (yeast extract-24 g/L, tryptone-12 g/L, glycerol-4 ml/L, KH_2_PO_4_-17 mM, K_2_HPO_4_-72 mM). The plasmids containing 147 bp α-satellite sequence and 145 bp 601 sequence were kind gifts from Prof. Curtis Alexander Davey. The pUC 147 bp DNA used as competitor DNA in previous nucleosome array works (33) was cloned from the pUC vector. A plasmid containing 10 copies of this 147 bp pUC DNA was cloned into a pUC57 vector. The α-satellite, 601 and pUC DNA plasmids were grown at 37°C for 18 hours in TB medium. The plasmids were purified as described (34). Plasmid concentrations were adjusted to 2 mg/ml and digested with 75 units of enzyme/mg of the plasmid during overnight incubation at 37°C. The mononucleosome DNA fragment and vector were separated by PEG 6000 size fractionation in the presence of 10 mM MgCl_2_. The supernatant containing the blunt-ended DNA fragment was precipitated with a 3 times volume of ethanol then resuspended in TE buffer (10 mM Tris pH 7.5, 0.1 mM EDTA) before further purification by ion-exchange on a MonoQ 5/50 Gl column (GE Healthcare Pte Ltd Singapore) to remove traces of vector and recombination products. 2 mg of the sample was loaded onto the column at a flow rate of 0.5 ml/min followed by washing with 10 column volumes of buffer A (300 mM LiCl, 20 mM Tris pH 7.5, 1 mM EDTA) to remove remnant PEG 6000. The DNA was eluted employing a linear gradient of 40% to 65% of buffer B (750 mM LiCl, 20 mM Tris pH 7.5, 1 mM EDTA) across 55 column volumes. The resulting fractions underwent separation by 10% PAGE in 1X TBE. Fractions containing the monomeric DNA were combined, concentrated and buffer exchanged to TE (10 mM Tris pH 7.5, 0.1 mM EDTA) using a 10,000 MWCO concentrator (Merck Pte. Ltd, Singapore).

Histone expression and refolding into octamer was carried out employing established protocol (35) and as described earlier (36) using pET3a plasmids (a kind gift from Prof. Curtis Alexander Davey) containing core human histones.

### Nucleosome reconstitution

The components for reconstitution were set up in a slide-A-dialyzer (volume < 1 mL) (Thermo Fisher Singapore Pte Ltd.) or dialysis bag (volume > 1 ml) (10 kDa MWCO). They were mixed in the following order: autoclaved water, then solutions 1 M Tris pH 7.5, 4 M LiCl, 0.5 M EDTA, 0.5 M DTT, human histone octamer (hHO) and last DNA. The final concentrations of the components in the dialysis mixture are 20 mM Tris pH 7.5, 2 M LiCl, 1 mM EDTA, 1 mM DTT, 4.8-7.2 μM hHO and 6 μM DNA. The dialysis mixture was then equilibrated against high salt buffer 20 mM Tris pH 7.5, 2 M LiCl, 1 mM EDTA and 1 mM DTT for 30 minutes in a beaker at room temperature. Following equilibration, the low salt buffer (20 mM Tris pH 7.5, 1 mM EDTA and 1 mM DTT) was slowly pumped into the dialysis set up under continuous stirring. While the low salt buffer was being pumped in, an equal volume of a mixture of high salt and the low salt buffer was withdrawn from the dialysis set up. Small-scale reconstitution was carried out by pumping 500 mL of the low salt buffer into 200 ml of high salt buffer, followed by a 4-hour incubation in the low salt buffer. In large-scale reconstitutions, 1500 mL of the low-salt buffer was continuously pumped into 600 ml of a high salt buffer across 18 hours, followed by a 4-hour incubation in the low salt buffer. Histone octamer saturation and formation of a well-defined NCP was observed as a single sharp band in the electrophoretic mobility shift assay (EMSA).

### Crystallization of the telomeric nucleosome

The telomeric DNA NCP was crystallised by employing an established protocol (37). The reconstituted NCP was prepared in 20 mM potassium cacodylate pH 6.0, 1 mM EDTA, 1 mM DTT and concentrated to a final concentration of 8 mg/mL. Telomeric DNA NCP crystals were grown as hanging drops comprising an equal volume of Telo-NCP and precipitant solution (63-68 mM MnCl_2_, 96-106 mM KCl, 20 mM potassium cacodylate pH 6.0). Crystals were harvested to the well containing 25 mM MnCl_2_, 30 mM KCl, 10 mM potassium cacodylate pH 6.0. The crystals were buffer exchanged to a final cryoprotectant solution (25 mM MnCl_2_, 30 mM KCl, 10 mM potassium cacodylate pH 6.0, 24% MPD, 5% trehalose) through 8 steps with a 10-minute incubation at each step and overnight incubation in the final buffer. The crystals were then cooled to 100 K in a liquid nitrogen stream followed by flash freezing in liquid nitrogen.

### Collection of X-ray diffraction data from native crystals

Diffraction intensities were collected at the X06DA beamline equipped with the PILATUS 2M-F detector at the Swiss Light Source (Villigen PSI, Switzerland), to a resolution of 2.6 Å for Dataset-1 and to a resolution of 2.2 Å for Dataset-2. Data collection was carried out at 100 K for 0-360° at wavelengths 2.072 and 1.000 Å with the oscillation of 0.25° and with an exposure of 0.2 seconds. Integration, scaling, and merging of the intensities was carried out using AutoProc (38,39).

### Data collection of Single-wavelength anomalous maps

All phased anomalous difference Fourier maps were collected at the beamline X06DA (PXIII) at (Swiss Light Source (SLS), Villigen PSI, Switzerland. Multi-orientation data collection was carried out at various χ (5°,10°,15°,20°,25°,30°,35°) and φ (60°, 90°, 120°) settings of the multi-axis PRIGo goniometer (40) and at a wavelength of 2.075 Å (phosphorus absorption edge), 1.8961 Å (manganese absorption edge) and 1.6038 Å (cobalt absorption edge), on a PILATUS 2M-F detector (41). The data were processed using XDS, scaled and merged using XSCALE (42) and phased by SHELX C/D/E (43) with anomalous peak heights calculated using AnoDe without a resolution cutoff (44). Refinement and model map cross-correlation calculations were performed using Refmac (45). Structure figures were prepared and rendered using Pymol (46).

Only for the cobalt dataset, the crystals were buffer exchanged to a final cryoprotectant (25 mM CoCl_2_, 30 mM KCl, 10 mM potassium cacodylate pH 6.0, 24% MPD) after cryoprotection. The buffer exchange was repeated 10 times, followed by overnight incubation, to remove all Mn^2+^.

### Structure determination of the telomeric nucleosome

Telomeric DNA nucleosome structures from Dataset-1, were determined by using Molrep (47) and 145 bp 601 NCP from *X. laevis* (PDB: 3LZ0 (28)) was used as a search probe. The structures were rebuilt and refined with the softwares COOT (48) and BUSTER (49). Two structures, hereafter named as Telo-NCP-1 (Orientation 1) and Telo-NCP-2 (Orientation 2), corresponding to the two orientations were obtained. Phases calculated from this model were subsequently used to locate phosphorus, manganese and cobalt sites using an anomalous Fourier map to ∼4.0 Å resolution at the phosphorus, manganese and cobalt absorption edge (50) (Supplementary Table S1). With Dataset-2, another telomeric nucleosome structure, hereafter named as Telo-NCP, was determined by using Molrep with Telo-NCP-1 as a search probe and refined with BUSTER (49). All structural graphics were generated by Pymol (46). The refined coordinates and structure factor amplitudes have been deposited to the Protein Data Bank with the PDB accession code 6L9H for the Telo-NCP-1, 6LE9 for the Telo-NCP-2 and 6KE9 for Telo-NCP.

### Tyrosine fluorescence measurements of NCP

Samples were prepared in 600 μl microfuge tubes by adding stock solutions in the order NaCl (4 M), Tris (1 M), EDTA (0.5 M), DTT (0.5 M) and finally NCP (10 μM) to a final concentration of 20 mM Tris, 1 mM EDTA, 1 mM DTT, 2 μM NCP and NaCl 0.2-3 M. The sample was equilibrated for one hour. Tyrosine fluorescence measurements were carried out in triplicate at 20 °C using a quartz cuvette (Hellma Asia Pte. Ltd. Singapore) at 1 cm path length on a Varian Cary Eclipse (Agilent Technologies Singapore Pte. Ltd) fluorescence spectrophotometer with excitation and emission wavelengths of 275 nm and 305 nm respectively. The measurements were normalised, plotted with Microsoft excel and fitted with Origin (Origin Lab, Northampton, MA) to derive the dissociation point (salt concentration at which 50% of nucleosome dissociated).

### Competitive mononucleosome reconstitution

Competitive reconstitution with salt dialysis and salt dialysis supplemented with 1 mM MgCl_2_ were carried out as for the NCP reconstitution (see above). The competitive reconstitution by salt dialysis and salt dialysis supplemented with MgCl_2_ made use of 200 ng ^32^P labelled tracer DNA of interest, 10 μg of calf-thymus core length DNA (145-147 bp mixed length obtained from the digestion of calf thymus chromatin), 5 μg of hHO in a 50 μl system. They were mixed in the following order: autoclaved water, then solutions 1 M Tris pH 7.5, 4 M LiCl, 0.5 M EDTA, 0.5 M DTT, hHO, calf-thymus core length DNA and last tracer DNA. The final concentrations of the components in the dialysis mixture are 20 mM Tris pH 7.5, 2 M LiCl, 1 mM EDTA, 1 mM DTT, 1.2 μM hHO, 2.4 μM calf-thymus core length DNA and 0.048 μM tracer DNA. Then small-scale nucleosome reconstitution was carried out as described above. For competitive reconstitution employing MgCl_2_, EDTA was eliminated from both high salt and low salt buffer and was supplemented with 1 mM MgCl_2_.

Nucleosome assembly in the presence of Nap1 and ACF was carried out using a Chromatin assembly kit (C01030001) (Diagenode Inc. United States), following the manufacturer’s protocol with minor modifications to accommodate tracer DNA. The reaction was carried out in a 25 μL aliquot containing 60 ng of ^32^P labelled tracer DNA of interest, 3 μg of calf-thymus core length DNA, 1.5 μg of hHO, 0.75 μL ATP (0.1M), 2.5 μL NAP1 (2 mg/mL), 2.5 μL ACF (0.2 mg/mL) and 2.5 μL of the 10X assembly buffer.

The reconstituted NCP was resolved by 6% PAGE using 0.2 X TB, followed by drying and exposure to image screen overnight. The visualisation was carried out using the typhoon FLA 700 system and the bands were quantified by Image J (51).

### Small-angle X-ray scattering (SAXS)

Solution samples were prepared by combining an appropriate volume of NCP stock with the buffer in a 600 μL microfuge tube and mixing by gentle flicking. The sample was transferred to 1.5-2 mm quartz capillaries (Charles Supper Company, USA) and sealed with wax.

Measurements were carried out at the BL23A SWAXS end station at the National Synchrotron Radiation Research Center, Hsinchu, Taiwan. Data were collected at a wavelength of 0.886 Å, using a CCD (charged coupled device) detector (MarCCD165, Mar Evanston, IL, USA) for 100-300 seconds at room temperature (25°C) at a sample-to-detector distance of 1.75 m. The data were corrected for capillary and solvent scattering and analyzed by PRIMUS and GNOM from the ATSAS package (version 2.8.2) (52,53). The setting of this software adheres to quality checks defined in the recent guidelines for publication of SAXS data (54). The respective subroutines of the ATSAS program perform a search for the optimal data range in determination of the radius of gyration R_g_ by the Guinier method and from the P(r) distribution; as well as in evaluation of particle diameter (D_max_) and in the fitting of theoretical form factors to the experimental SAXS profiles.

## RESULTS AND DISCUSSION

### Crystal structure of the telomeric NCP

To understand how human telomeric DNA forms nucleosomes and chromatin, we set out to reconstitute NCPs from a 145 bp telomeric DNA sequence with the aim of determining the crystal structure. As human telomeric DNA has the propensity to form G-quadruplexes in the presence of Na^+^ or K^+^ (55), the standard nucleosome salt reconstitution protocol (34) was modified substituting NaCl for the same concentration of LiCl. This resulted in the formation of a well-defined NCP characterised by the appearance of a single sharp band in the EMSA gel (Supplementary Figure S1). We employed similar crystallisation conditions as previously published (37,56) and obtained hexagonal rod crystals in the presence of MnCl_2_, characteristic of NCP crystals.

The 145 bp telomeric DNA NCP structure was determined by molecular replacement using 3LZ0 (28) as a search probe, followed by collecting anomalous peaks from the phosphorus edge to identify the coordinates of the 114 phosphorous atoms on the telomeric DNA backbone (Supplementary Figure S2a). The resulting electron density map showed diffuse densities (data not shown) on the DNA bases, which indicated an average of two NCP orientations. Manganese and cobalt anomalous maps were used to verify that this was the case (Supplementary Table S1). A total of 15 datasets were collected from a single crystal diffracting at ∼3.0 Å resolution in multiple orientations at a wavelength of 1.8961 Å and the datasets were merged to locate manganese positions from the anomalous peaks. In addition, 7 datasets were collected at a wavelength of 1.6083 Å to locate cobalt position from the anomalous peak. Whilst Mn^2+^ and Co^2+^ ions are known to coordinate with N7 and O6 of guanine in ‘GG’ or ‘GC’ steps located on one of the DNA strands in telomeric DNA (57,58), the anomalous data showed that both Mn^2+^ and Co^2+^ were distributed on both DNA strands (Supplementary Figure S2b,c and S3a). Since, this is chemically impossible for the telomeric DNA sequence, our observation instead indicated a crystal in which the NCP packs in two orientations as discussed in (59). In agreement, we saw the best fit by refining this dataset for a crystal comprising two orientations in equal proportions, diffracting anisotropically at a resolution of 2.5 Å (called Dataset-1) (Table 1). From this, we generated two separate PDB files corresponding to the two orientations: Telo-NCP-1 (Orientation-1, PDB code 6L9H) (Figure 1a) and Telo-NCP-2 (Orientation-2, PDB code 6LE9) (Figure 1b). We observed similar Mn^2+^ distributions for the two orientations and noted two modes of coordination with N7 of guanine (Figure 1c and d), and comparable Co^2+^ binding (Supplementary Figure S3e). Two orientations of the NCP in the crystal lattice has been observed previously for the 145 bp non-palindromic 601 DNA sequences with *X. laevis* histones (PDB codes 3LZ0, 3LZ1 (28)).

**Table 1.** Data statistics for the telomeric NCP.

| Data collection and Processing | Dataset-1<br>(Two orientations) | Dataset-2<br>(Single orientation) |  |
| --- | --- | --- | --- |
| X-ray source | X06DA (SLS) | X06DA (SLS) |  |
| Wavelength (Å) | 2.072 | 1.000 |  |
| Space group | P2<sub>1</sub>2<sub>1</sub>2<sub>1</sub> | P2<sub>1</sub>2<sub>1</sub>2<sub>1</sub> |  |
| Unit-cell dimensions |  |  |  |
| a, b, c (Å) | 106.45, 109.38, 176.37 | 106.13, 109.32, 176.97 |  |
| α, β, γ (°) | 90.0, 90.0, 90.0 | 90.0, 90.0, 90.0 |  |
| *Isotropic* |  |  |  |
| Resolution range<sup>a</sup> (Å) | 92.96 – 3.29 (3.35 – 3.29) | 93.01 – 2.97 (3.02 – 2.97) |  |
| R<sub>merge</sub><sup>a</sup> | 0.112 (1.305) | 0.048 (0.988) |  |
| <I/σ(I)><sup>a</sup> | 14.5 (2.2) | 26.7 (2.3) |  |
| CC<sub>1/2</sub><sup>a</sup> | 0.997 (0.947) | 1.000 (0.973) |  |
| R<sub>pim</sub> | 0.03 (0.37) | 0.01 (0.27) |  |
| Completeness (%) | 99.9 (99.7) | 99.8 (100.0) |  |
| Redundancy<sup>a</sup> | 13.0 (13.4) | 13.4 (14.3) |  |
| *Anisotropic* |  |  |  |
| Resolution range<sup>a</sup> (Å) | 92.96 – 2.52 (2.89 – 2.52) | 93.0 – 2.18 (2.55 – 2.18) |  |
| Ellipsoidal resolution<sup>b</sup> (Å) / direction | 3.53 / *a* * | 3.21 / *a* * |  |
|  | 2.49 / *b* * | 2.18 / *b* * |  |
|  | 3.68 / *c* * | 3.38 / *c* * |  |
| R<sub>merge</sub><sup>a</sup> | 0.114 (1.506) | 0.050 (1.231) |  |
| <I/σ(I)><sup>a</sup> | 14.3 (1.9) | 26.1 (2.1) |  |
| CC<sub>1/2</sub><sup>a</sup> | 0.998 (0.795) | 1.000 (0.808) |  |
| R<sub>pim</sub><sup>a</sup> | 0.03 (0.43) | 0.01 (0.34) |  |
| Completeness, spherical<sup>a</sup> (%) | 47.3 (7.0) | 42.1 (5.7) |  |
| Completeness, ellipsoidal<sup>a,b</sup> (%) | 94.5 (79.6) | 94.0 (81.2) |  |
| Redundancy<sup>a</sup> | 13.0 | 13.8 |  |
| **Structure Refinement** |  |  |  |
|  | **Telo-NCP-1** | **Telo-NCP-2** | **Telo-NCP** |
| PDB code | 6L9H | 6LE9 | 6KE9 |
| Resolution range (Å) | 49.00 – 2.60 | 49.00 – 2.60 | 42.01 – 2.22 |
| Refinement (R<sub>work</sub>/R<sub>free</sub>) | 23.8/27.8 | 24.1/28.1 | 20.5/23.0 |
| r.m.s.d distances (Å<sup>2</sup>) | 0.007 | 0.007 | 0.010 |
| r.m.s.d bond angles (°) | 0.76 | 0.77 | 0.94 |
| Estimated coordinate error (Å) | 0.000 | 0.000 | 0.000 |
| *Ramachandran plot* |  |  |  |
| Favored regions (%) | 95.25 | 94.44 | 96.12 |
| Allowed regions (%) | 4.75 | 5.56 | 3.88 |
| Disallowed regions (%) | 0.00 | 0.00 | 0.00 |

<sup>a</sup>Values in parentheses are for the highest-resolution shell.<sup>b</sup>The datasets were anisotropically truncated using the STARANISO. The anisotropic cut-off surface was fitted to an ellipsoid to provide an approximate resolution limit in the three directions in reciprocal space. The real cut-off surface is only approximately ellipsoidal, and the directions of the worst and best resolution limits may not correspond with the reciprocal axes.

**Figure 1.**
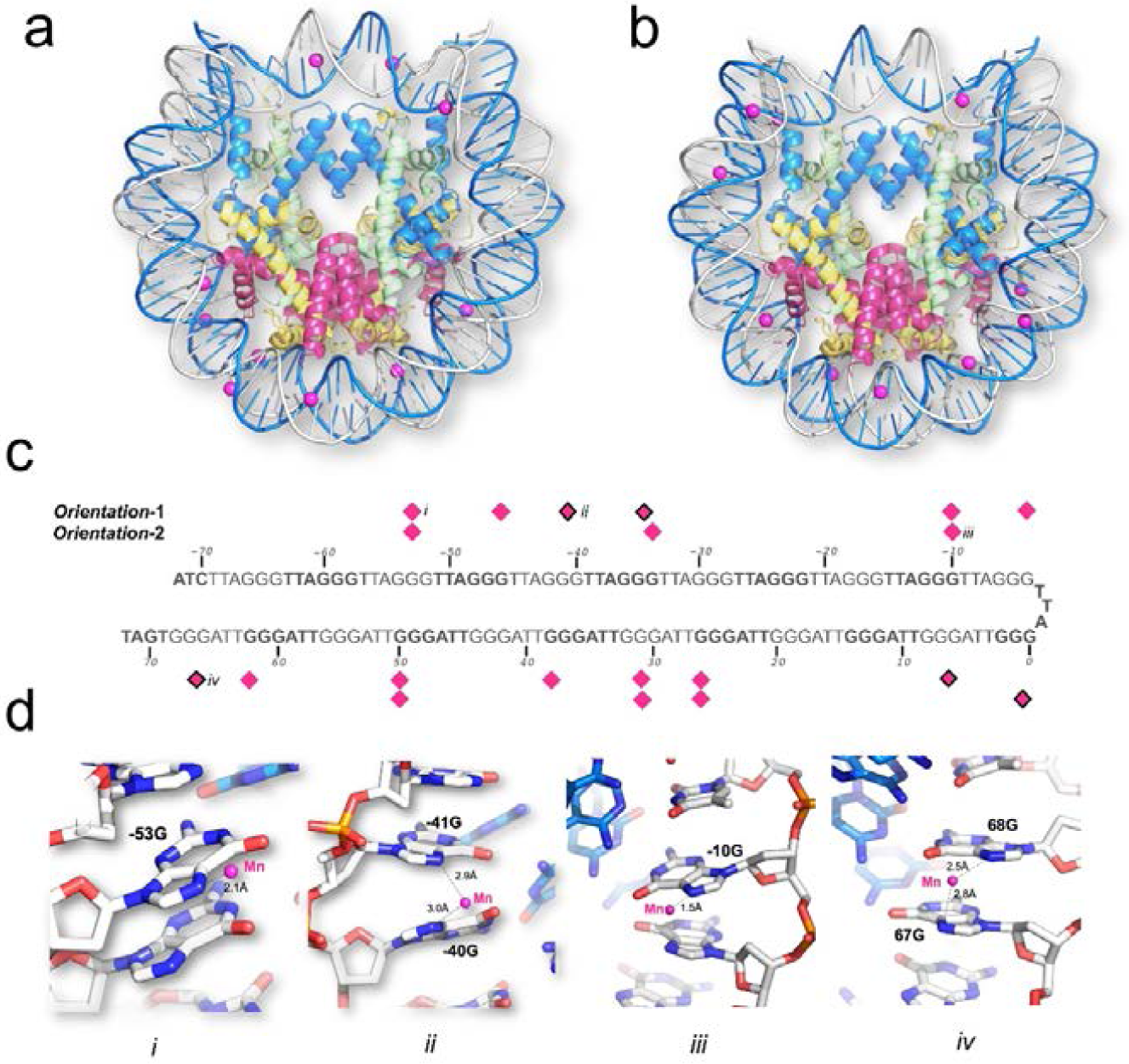
The two-orientation telomeric NCP crystal structure solved by a combination of molecular replacement, and single-wavelength anomalous maps at the phosphorous, manganese and cobalt absorption edge. Mn^2+^ coordination to the N7 guanine atoms in (**a**) Telo-NCP-1 (Orientation-1) and (**b**) Telo-NCP-2 (Orientation-2). (**c**) Coordination pattern of the Mn^2+^ atoms in Telo-NCP-1 and Telo-NCP-2. The Mn^2+^ shows two modes of coordination: Mn^2+^ bound to a single N7 guanine (shown as magenta diamonds), and Mn^2+^ interacting with N7 of two adjacent guanine bases (marked as magenta diamonds with black borders). (**d**) Examples of Mn^2+^ interaction with DNA are shown in panels ***i ii, iii*** and ***iv***. Panels ***i*** and ***iii*** show Mn^2+^ coordination to a single N7 guanine in Telo-NCP-1 and Telo-NCP-2 respectively. Panels ***ii*** and ***iv*** display Mn^2+^ interaction with N7 of two adjacent guanine bases in the Telo-NCP-1 and Telo-NCP-2 respectively. Henceforth, for illustration, the most frequent naming of DNA and proteins chains is adopted. The two DNA strands are named as ‘I’ and ‘J’, with nucleotides on each strand numbered from −72 bp to 72 bp. The nucleotides are labelled with a numeral identifier followed by strand identifier, e.g. ‘−2I’ corresponding to nucleotide number ‘−2’ from the strand I and histones are labelled A and E (H3) (blue); B and F (H4) (green); C and G (H2A) (yellow); and D and H (H2B) (red).

By screening several crystals, we identified one that gave excellent diffraction and hypothesised that this comprised NCPs with a single orientation in the crystal lattice. We confirmed this by using Telo-NCP-1 and Telo-NCP-2 structures for molecular replacement, followed by refinement with BUSTER (49). The refinement statistics from Dataset-2 showed a clear preference for orientation-1 (Supplementary Table S2). We confirmed this observation by generating omit maps at −28I guanine. Substitution guanine with other DNA bases (adenine, thymine and cytosine) resulted in negative peaks in Fo−Fc map (Supplementary Fig. S4) and the Rfree values showed a preference for guanine (Supplementary Table S2). We did not collect an anomalous data set for this crystal. However, putative manganese densities at the coordination distance from the N7 of guanine were observed in the Fo−Fc map. The Telo-NCP-1 showed a higher number of putative manganese densities in comparison with Telo-NCP-2 (8 versus 4) (Supplementary Figs. S5 and S6). Preference for a single orientation has been observed previously for an NCP with a non-palindromic DNA sequence containing a poly (dA·dT) tract as well as for the MMTV-A NCP (29,59). In the Telo-NCP refined map, the electron density showed well-resolved bases as well as distinct densities characterising purines and pyrimidines (Supplementary Figure S7).

The crystal packing and general architecture of Telo-NCP at 2.2 Å resolution (Figure 2a) are similar to that of other published NCP crystal structures (60). The HO protein structure in the Telo-NCP showed global superimposed RMSD deviations of 0.30 and 0.29 Å, compared to the histone fold regions of the 145 bp α-satellite *X. laevis* NCP (PDB code 2NZD (61)) and 145 bp 601 *X. laevis* NCP (PDB code 3LZ0 (28)). We also observed the DNA “stretching” around super-helical locations (SHL) ± 5, required for the DNA ends to pack within the crystal lattice. The precise location of such stretching, however, influences the positioning of flexible base steps in the inward locations of the narrow minor groove at the so-called ‘pressure points’, where the phosphates ‘hook’ the H3/H4 histones (Figure 2b) (62,63). In the Telo-NCP structure, there are several flexible TA and TT base steps located in the regions of inward-facing minor grooves (Figure 2b and 2c).

**Figure 2.**
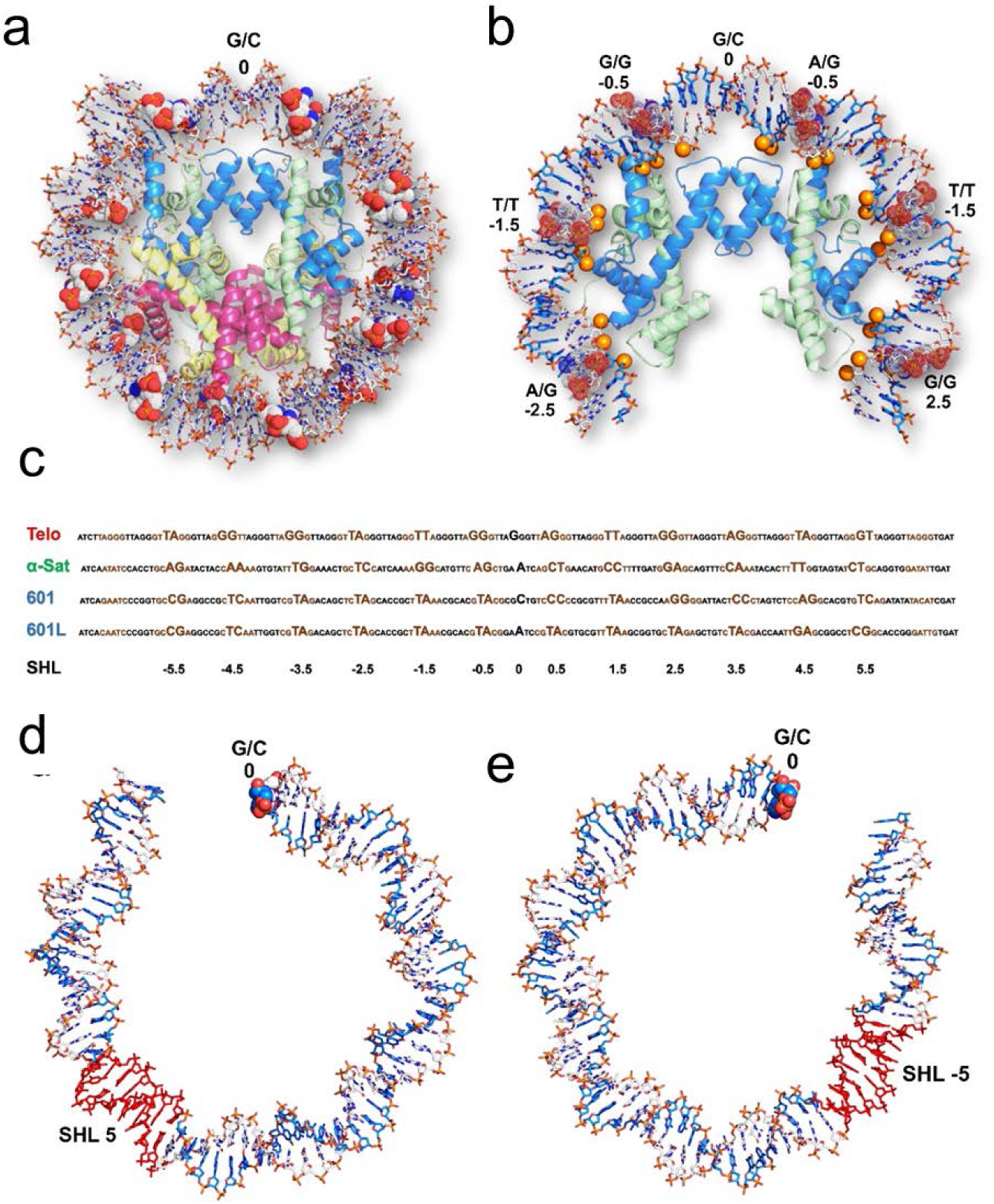
Overall structure of the 2.2 Å-resolution telomeric NCP. (a) Telo-NCP structure with DNA shown in stick and histone octamer in cartoon representation. The DNA is colored by elements and the four histones H3, H4, H2A and H2B are colored blue, green, yellow and red respectively. The base steps at minor groove pressure points are shown as space filling dots. (**b**) Base steps between SHL −2.5 to SHL 2.5 at the minor groove pressure points in Telo-NCP. The DNA phosphates that mediate histone-DNA interaction are shown as orange spheres. (**c**) Comparison of the positioning of the telomeric DNA, α-satellite, 601 and 601L DNA on the histone octamer. The major groove elements are shown in black and minor groove elements in brown. The base steps at the minor grove facing the histone octamer and dyad position are highlighted. **(d)** The stretching (red) on the telomeric NCP between SHL 4.5 to 5. **(e)** The stretching (red) on the telomeric NCP between SHL −4.5 to −5.

### Flexible base steps in the telomeric NCP enables the formation of a well-defined DNA path on the HO

To understand how the telomeric DNA sequence wraps around the HO making the appropriate contacts to form an NCP, we compared the DNA path as seen in the 2.2 Å single-orientation Telo-NCP structure with that of the 2.5 Å 145 bp ‘601’ (PDB code 3LZ0 (28)) and the 2.6 Å α-satellite NCPs (PDB code 2NZD (61)). Remarkably, the Telo-NCP DNA stretching at SHL ±5 (Figure 2d and 2e) occurs in an almost identical location in the 601-NCP, resulting in similar DNA paths and phosphate locations (Figure 3a). By contrast, when we compared phosphate positions with the 145 bp α-satellite NCP, the Telo-NCP had two regions with major deviations of around 6Å at bp locations −45I to −20I and 28I to 50I (Figure 3b). We deduced that these large deviations resulted from the location of the symmetric DNA stretching at the two SHL ±5 in Telo-NCP, which in α-sat NCP is located at SHL ±2. Therefore, the DNA in Telo-NCP is out of phase by one base pair from 28I to 50I (SHL 2.5 to SHL 5) compared to the corresponding phosphates of the α-satellite NCP (Supplementary Figure S8).

**Figure 3.**
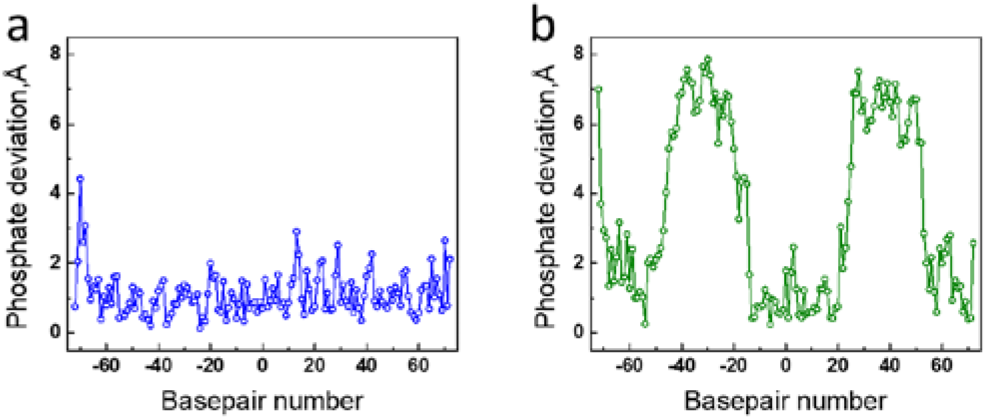
Differences in the DNA path in the crystals of the 2.2 Å-resolution Telo-NCP in comparison with 145 bp NCP with the 601 sequence and α-satellite sequence (PDB codes 3LZ0 and 2NZD respectively). Distances between DNA phosphorous atom locations between superimposed structures of the Telo-NCP and respective NCPs are shown as a function of base location in the I DNA strand. (**a**) Phosphate deviations between Telo-NCP and 601-NCP. (**b**) Phosphate deviations between Telo-NCP and α-satellite NCP.

In order to reveal how the telomeric DNA distorts to form an NCP, we analysed the minor and major groove inward-facing regions, as well as the locations of the so-called ‘pressure points’; the base steps where the minor groove most closely faces the HO (62,63). These pressure points are defined by the base step locations where the (H3/H4)_2_ tetramer binds to DNA around the nucleosome dyad and hook the DNA phosphates by polar contacts at the minor groove inward-facing registers (62,63). We aligned the Telo-NCP DNA sequence with the 145 bp 601, 601L and α-satellite NCPs (Figure 2c and Supplementary Table S3). In the 145 bp Telo-NCP DNA there are 23 TT, TA, AG, and GT steps as well as the 46 ‘GG’ steps. It is known from analyses of DNA structures that among these base steps, the most flexible is the TA step, followed by AG, TT, GG and finally GT, with corresponding stacking energies of −0.19 kcal/mol (TA), −1.06 kcal/mol (AG), −1.11 kcal/mol (TT), −1.44 kcal/mol (GG) and −1.81 kcal/mol (GT) (64). Flexible base steps, in particular the TA step, are most able to adjust to the structural deformations necessary to accommodate the bending at the pressure point HO contact locations. Accordingly, we observed that in the Telo-NCP, 5 of the 12 central (around the dyad) minor groove inward pressure points contain TA or TT steps and two contain the AG steps (Figure 2c and Supplementary Table S3). Furthermore, the presence of several additional TA steps in both the minor and major groove inward-facing registers also appears to be important for bending the DNA to facilitate the wrapping that enables the phosphate interactions with the HO. The DNA distortions required for this bending result in several base-step parameters displaying considerable deviations from “ideal” values (notably roll and twist, see below). For comparison, the α-satellite NCP has eight favourable flexible base steps located in the 12 central minor groove inward facing positions around the dyad, but none of them is a TA step. The NCP structure of the highest known nucleosome affinity DNA sequence, the 145 bp 601L (62), has TA steps occupying 8 of the 12 pressure points in the minor groove, while the “601” has 5 such steps (Figure 2c and Supplementary Table S3). A second characteristic of importance for nucleosome positioning that is inherent in the Telo-NCP sequence, and which it shares with the ‘601’ sequence, is its high GC content (50%). As a result, in 8 out of the 12 central positions, GG steps are located in the wide major groove inward registers while we identified 4 GG steps at minor groove pressure point locations (Figure 2c). It may be noted that in the two-orientation 2.6 Å telomeric NCP structure, the location of all of the minor groove inward-facing pressure point base steps at the HO contact points, are the same as in the 2.2 Å structure (Supplementary Table S3). The two Telo-NCP structures also have the same stretching location, DNA path and almost identical overall structures.

### The telomeric NCP displays pronounced DNA deformations

Next, we analysed how the telomeric DNA structure deforms from its native state to permit wrapping around the HO and also compared it to the DNA structure in other NCP structures. The base-step parameters of Telo-NCP DNA were plotted and compared to those in the NCPs formed from the 145 bp palindromic ‘601L’ (NCP-601L, PDB code 3UT9 (62)) and α-satellite (PDB code 2NZD (61)) DNA sequences (Figure 4 and Supplementary Figure S9). We observed that many TA elements displayed highly pronounced deviations in base-step parameter values (relative to average B-DNA (65)), and distortions were significantly larger compared to those of the NCP-601L and α-satellite NCP. In particular, roll values showed considerable deviations but slide, tilt, and shift values also deviated significantly. Similarly, a large proportion of the other base steps in the Telo-NCP exhibited considerable deviations (Figure 4). These base step parameter deviations originated from pronounced structural deformations at inward minor groove pressure points as well as other locations (Supplementary Figure S10). Interestingly, we also found that several GG and GT steps showed large deviations from the ideal base step parameter values. For example, we noted a highly pronounced tilt of 36 degrees for the GT step 15 at SHL 1.5 (Supplementary Figures S9 and S10). This analysis reveals that a large number of flexible base-steps present in the telomeric DNA are able to deform considerably in order to accommodate the DNA bending, which in turn enables the contacts between the inward-facing minor groove phosphates of the DNA with the HO, required for NCP formation.

**Figure 4.**
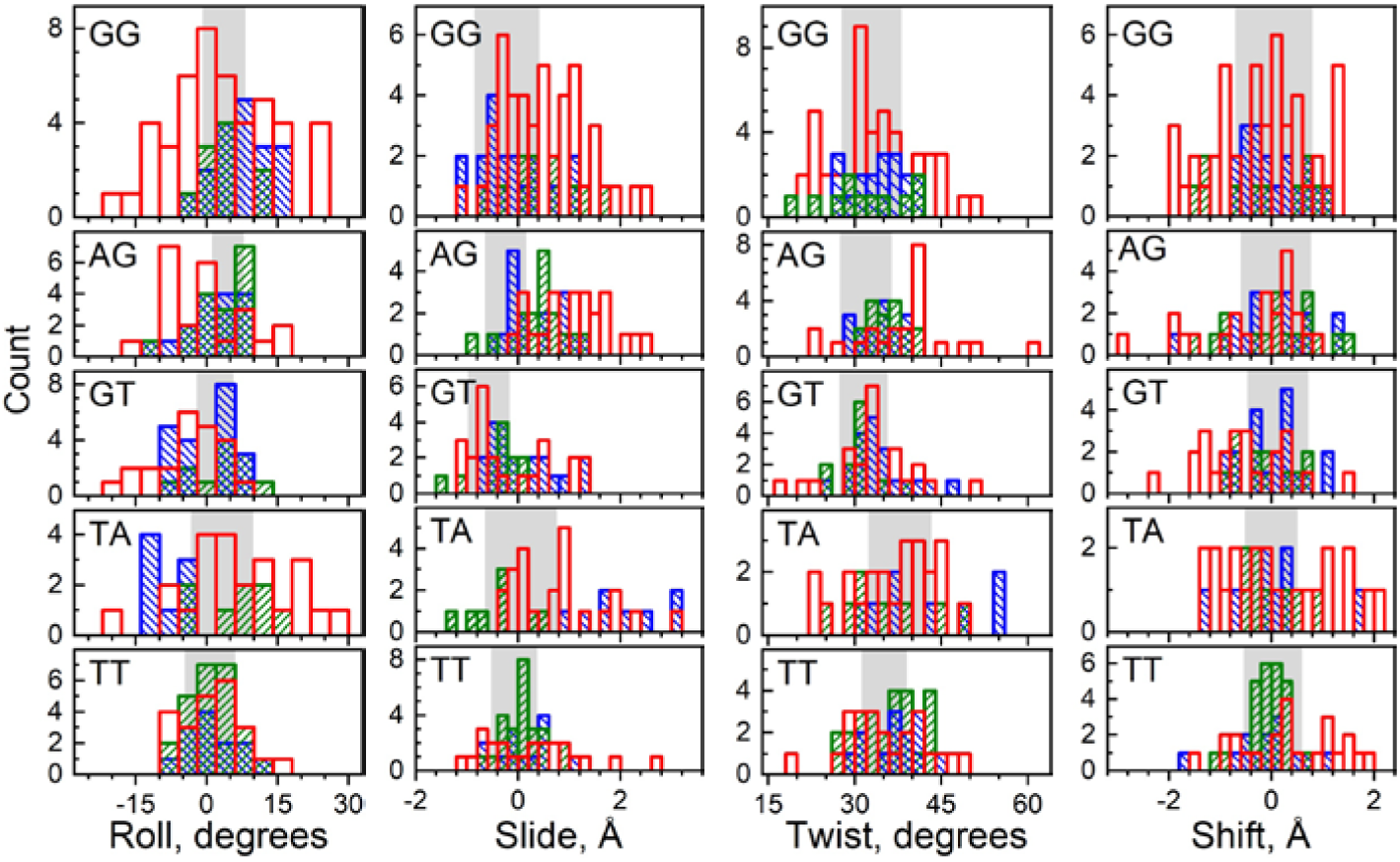
Population distribution of roll, slide, twist and shift base step parameters in Telo-NCP compared with the same steps present in NCP-601L and α-satellite NCP. A complete illustration of all six base step parameters (also including tilt and rise) including all base steps in NCP-601L and α-satellite NCP is shown in Supplementary Figure S9). Base steps from Telo-NCP, NCP-601L and α-satellite NCPs are shown respectively as red, blue and green columns. In each panel the shaded area covers the range of values (mean ± s.d.) observed in DNA crystals according to the analysis by Olson et al. (65). A majority of the base steps in the Telo-NCP and especially the TA and GG steps show significantly broader distribution and larger deviation from ideal B-DNA parameters compared to that observed for the NCP-601L and α-satellite NCP. See Supplementary Figure S10 for the illustrations of base steps showing significant distortions.

### The basic H4 N-terminal tail interacts with the HO acidic patch and stabilises nucleosome stacking in the crystal

In other NCP structures (27,36,66-70), the basic H4 tail 16-23 aa domain mediates nucleosome stacking by interacting with the acidic patch of H2A-H2B located on the surface of the neighbouring HO core and thereby regulates chromatin fibre compaction (27,36,66-70). We wondered whether this H4 tail interaction was also present in the Telo-NCP crystal structure, which would suggest a similar stabilisation of telomeric chromatin compaction by H4 tail-mediated nucleosome-nucleosome stacking. The structure of the 17-23 aa domain of one of the H4 histones (chain ‘B’) was well resolved and we found that the details of the interaction with the H2A-H2B acidic patch of the neighbouring NCP were present in the Telo-NCP crystal structure (Figure 5a). Arg17 from the H4 tail (chain ‘B’) also forms hydrogen bonds with the −21I DNA phosphate group of its own NCP (Figure 5b). By comparison, none of the Arg 17 aa of the H4 tails of the palindromic NCP-601L (62) are in contact with DNA (Figure 5c). We speculate that this strengthened tail interaction with DNA might contribute to the preference for a single orientation in the crystal lattice, as observed in the non-palindromic 147 bp MMTV-A NCP (29).

**Figure 5.**
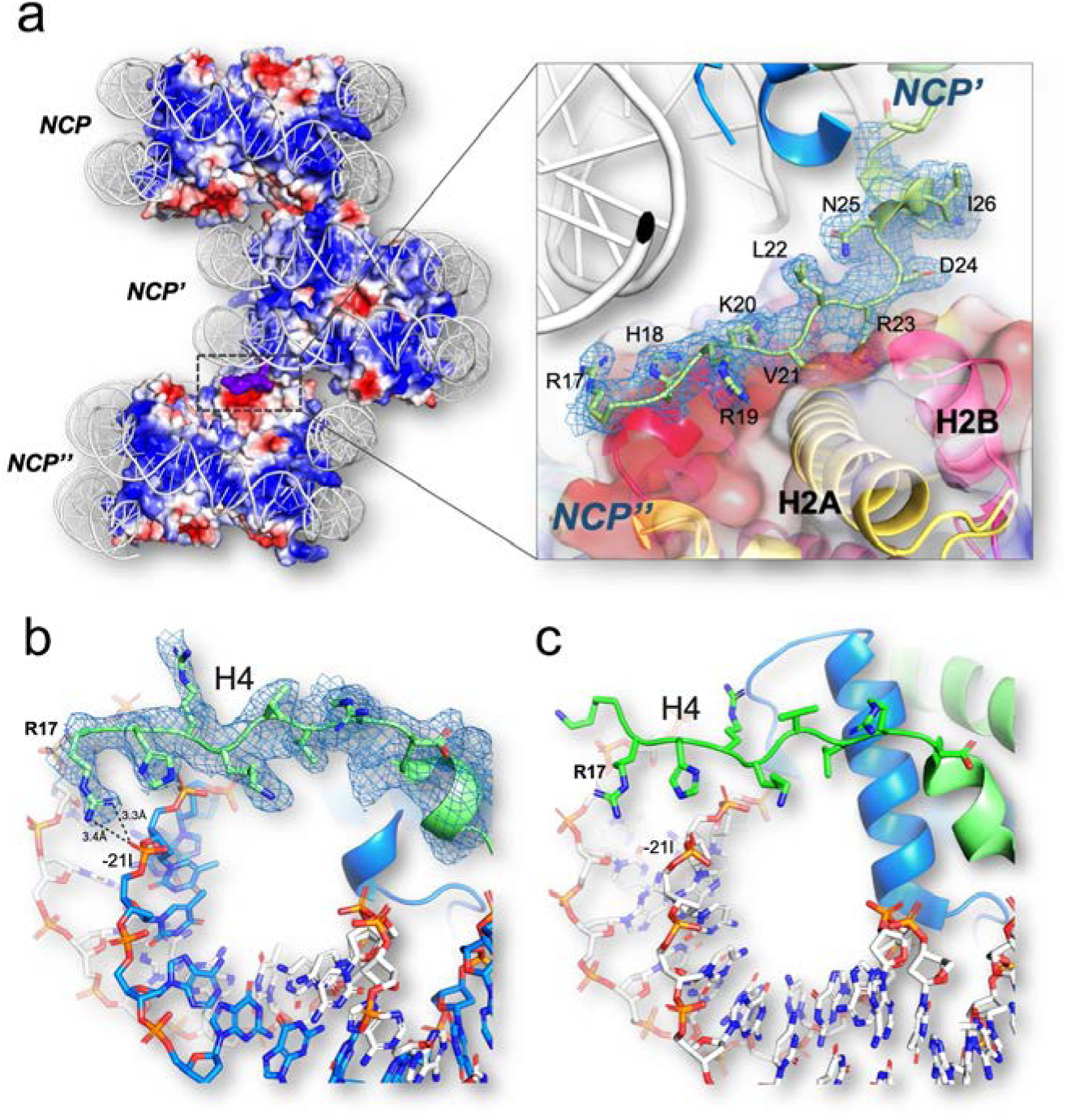
H4 tail interactions in Telo-NCP. **(a)** Illustration of the H4 tail mediated NCP-NCP stacking in the NCP crystal through interaction between the H4 Arg19-Arg23 amino acids and the H2A/H2B acidic patch of the adjacent NCP. Red and blue regions indicate domains of negative and positive electrostatic potential, respectively. The basic H4 tail is shown in purple. The zoom-in to the right shows that the H4 tail (chain ‘B’) has a well-defined electron density as observed from the 2Fo−Fc map contoured at 1.0 σ and it displays interaction with the HO core acidic patch. **(b)** The Arg17 of the H4 tail interacts with the phosphate of −21I (thymine). **(c)** The H4 tail in the NCP-601L does not interact with the DNA.

### The Telo-NCP exhibits a stable yet dynamic structure in solution

Telomeres are a hotspot for DNA damage and a target for the DNA damage response (DDR), but the molecular basis of their susceptibility is incompletely understood. We hypothesised that a contributing factor might be differences in stability and dynamic solution properties of Telo-NCP, compared to those of NCPs reconstituted with a nucleosome positioning sequence. We first carried out a salt-dependent nucleosome dissociation assay to compare the stability of Telo-NCP, the 145 bp 601 (601-NCP), the 147 bp α satellite (α-sat-NCP) and a random DNA sequence from the digested pUC vector backbone (pUC-NCP). The salt molarity at which 50% of the NCP dissociates defines its dissociation point (see Methods and Supplementary Figure S11). The dissociation point for Telo-NCP (1.04 M) was markedly lower than that of 601-NCP (1.53 M NaCl), and lower than that of α-sat-NCP (1.08 M NaCl) and pUC-NCP (1.13 M NaCl) (Figure 6a). This suggests that once formed, the assembled telomeric nucleosome is less stable than NCPs formed from other DNA sequences.

**Figure 6.**
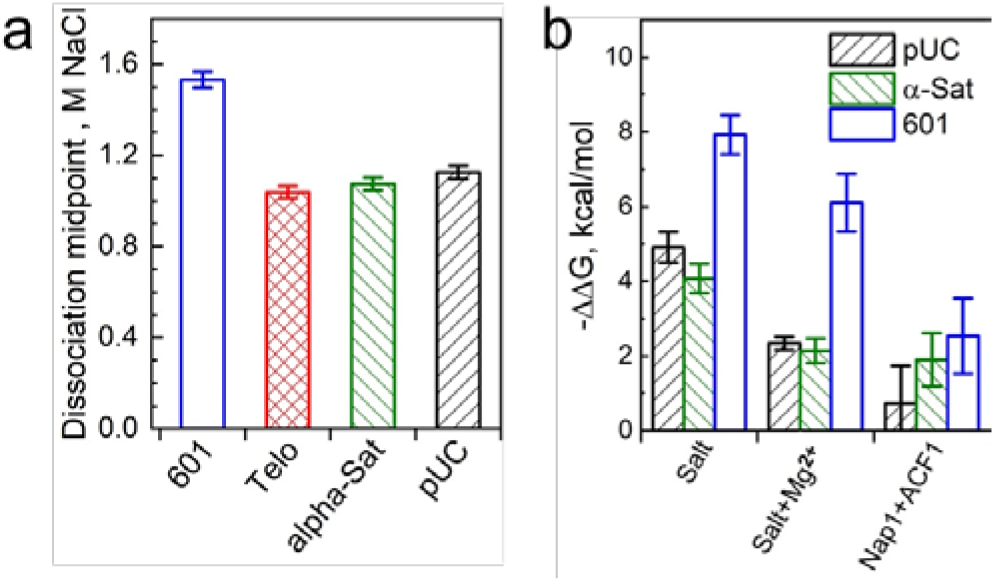
Stability of Telo-NCP and telomeric DNA affinity for the histone octamer compared to NCPs with various DNA sequences. (**a**) Salt-dependent stability of different NCPs in the NaCl buffer. The 601-NCP is shown in blue, Telo-NCP in red, α-Sat-NCP in green and pUC-NCP in black. The dissociation points of different NCPs were calculated from the curves of NaCl-induced NCP dissociation (shown in Supplementary Figure S11). **(b)** Relative affinity of the telomeric DNA for the HO. The difference in free energy values (ΔΔG) compared to the competitor calf-thymus core length DNA (see Methods for details) obtained for the various sequences were normalized using the results recorded for telomeric DNA as reference (ΔG = 0 kcal/mol). ΔΔG of NCP formation relative to Telo-NCP for the three DNA sequences is shown for three sets of competitive reconstitution experiments. 601-NCP is shown in blue, α-sat-NCP in green and pUC-NCP in black.

Next, we compared affinities of the telomeric DNA sequence for the hHO with those of the three other DNA sequences (601-NCP, α-sat NCP and the random pUC-NCP sequence) using a competitive reconstitution assay (Figure 6b, Supplementary Figure S12). We employed three different reconstitution conditions: salt dialysis, salt dialysis in the presence of 1 mM Mg^2+^, and in the presence of a mixture of the nucleosome assembly factor ACF and the chaperone Nap1. In agreement with earlier data (20), under standard salt dialysis conditions, telomeric DNA has the lowest affinity for HO among the tested DNA sequences (Figure 6b, Supplementary Figure S12a). The differences in free energy values (ΔΔG) of telomeric DNA compared to the other sequences were 7.9 kcal/mol for 601 DNA, 4 kcal/mol for α-satellite DNA and 4.9 kcal/mol for the pUC DNA (Figure 6b). In the presence of 1 mM Mg^2+,^ we observed reduced ΔΔG values compared to standard salt dialysis conditions. The NCP bands obtained for NCP assembly with the nucleosome assembly kit were unclear (Supplementary Fig. S12c). The fraction of DNA incorporated into the nucleosome was lower in comparison to that for reconstitution by salt dialysis. ΔΔG values of NCP formation relative to Telo-NCP were further reduced for the three DNA sequences in the presence of ACF and Nap1. Relative to α-sat NCP, the affinity difference is about 1.5RT, which suggests that under *in vivo*-like conditions, the affinity of the telomeric DNA for the human HO is somewhat lower than that of other natural positioning sequences.

We then went on to investigate the structural and dynamic properties of Telo-NCP in solution by small-angle X-ray scattering (SAXS), which gives information about averaged NCP size and DNA unwrapping at the ends. The experimental SAXS spectra at different concentrations were fitted in order to obtain the NCP particle radius of gyration (R_g_), distance distribution function, P(r), and maximal particle diameter (D_max_) (52). We saw that Telo-NCP (R_g_ = 46.3 Å) appeared larger than the 601-NCP (R_g_ = 42.1 Å) with D_max_ of Telo-NCP being 29 Å larger than that of 601-NCP (Supplementary Figure S13a, Table S4). This conclusion was also supported by dynamic light scattering investigations, which displayed a larger hydrodynamic diameter for Telo-NCP in comparison with 601-NCP (data not shown). Earlier works have shown that the depth of the dip in the NCP SAXS spectrum at 0.14 Å^-1^ is sensitive to DNA unwrapping (71-74). The NCP at low concentration (1-2.5 mg/mL) in low salt buffer (10 mM KCl) exhibit minimal inter-particle interactions, thereby facilitating the study of the form factor of the NCP SAXS spectrum (52). To compare experimental data with atomic NCP structures modelling different degrees of DNA unwrapping, we theoretically generated a number of SAXS profiles (form factors). We used the crystal structure of the 145 bp NCP with the 601 sequence (PDB code 3LZ0 (28)), the Telo-NCP determined in the present work, and the 601 NCP structure obtained after 150 ns molecular dynamics simulations of all-atom NCP. The form factors obtained from the crystal structures lacking coordinates of the histone tails fitted poorly with the experimental data (Supplementary Table S4). The structures derived from the MD simulations that included all the tails at the positions characteristic for the NCP solution (tails collapsing and attached to the DNA) fitted well with the experimental SAXS profiles (Supplementary Fig. 13b and Table S4. We calculated the SAXS spectra from atomic structures of the NCP generated with up to 10 bp unwrapped from either or both ends of DNA and fitted them to the experimental spectra (Figure 7, Supplementary Figure S13c and Supplementary Table S4). The shallow minimum at 0.14 Å^−1^ for Telo NCP indicated unwrapping of DNA ends resulting in the larger R_g_ and D_max_ values. An NCP particle having up to 30 – 40 bp DNA asymmetrically unwrapped from one of the ends gave the best fit to the spectrum (Figure 7, Supplementary Figure S13c), while structures with symmetric DNA unwrapping did not fit well (Supplementary Table S4). This suggests that in solution, the Telo-NCP displays a distribution of HO positioning on the DNA with a preference for asymmetric unwrapping. On the contrary, we observed that the 601-NCP exhibited a central placement of the DNA on the HO.

**Figure 7.**
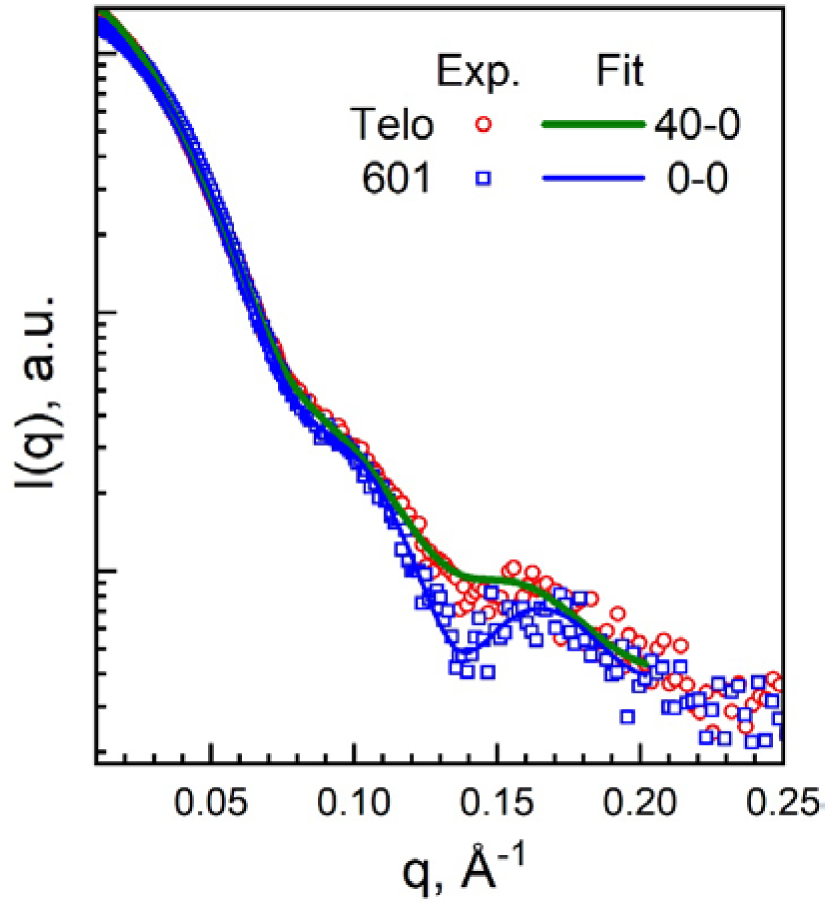
SAXS results for Telo- and 601-NCPs at low NCP concentrations where interparticle interactions are negligible. Experimental SAXS spectra of Telo- (red circles) and 601-NCP (blue squares) at 1 mg/mL NCP, 10 mM KCl, 10 mM Tris, pH 7.5, 1 mM EDTA, 1 mM DTT. Green (for the Telo-NCP) and blue (for the 601-NCP) curves are best-fits of the respective experimental spectra to the scattering profiles calculated for atomistic NCP structures modelling different degrees of DNA unwrapping from the histone core. See Methods, Supplementary Figure S13 and Supplementary Table S4 for more details of the fitting procedure.

The NCP reconstitution assisted by ACF and Nap1 represents a condition closer to that *in vivo*. Our results from the competitive reconstitution experiments, therefore, suggest that formation of the telomeric nucleosome and chromatin under *in vivo* conditions is not constrained by the inherent affinity of the HO for the telomeric DNA. Furthermore, for the three natural sequences tested, the difference in salt-dependent stability is less than 100 mM Na^+^, implying that the assembled telomeric nucleosome is rather stable, albeit less stable than other natural DNA sequences including natural positioning sequences.

We observed that the anomalous phosphorous peak was absent at one of the two ends of the DNA in the two orientation 2.6 Å structure, (Supplementary Figure S2a). The B-factors of this DNA end were also comparatively higher than those at the other end (Supplementary Figure S2a). Telo-NCP has two GG steps at pressure points in SHL −3.5, −4.5 compared to AG and TA steps at SHL 3.5 and 4.5 respectively. Unwrapping associated with these GG base step, which has paid a larger energy cost (compared to AG and TA steps) for the bending associated with the minor groove inward contact, likely leads to a preference for unwrapping at this end. This suggests that the precise location of these two GG steps at one end leads to the asymmetric unwrapping of 30-40 bp on the Telo NCP, which was inferred by the SAXS data. The asymmetric DNA unwrapping on the nucleosome core observed in the solution SAXS experiment is dependent on local DNA sequence (75,76) and has been observed previously (73,76,77). We also observed lower salt-dependent stability and lower affinity for Telo-NCP compared to the other DNA sequences investigated. All these features result from the unique repetitive telomeric DNA sequence, which gives the Telo-NCP its highly dynamic properties. These dynamic features may be important in facilitating the biological function and properties of telomers by mediating DDR, rapid chromatin remodelling and dynamic regulation of epigenetic states, thereby maintaining genomic integrity.

## CONCLUSIONS

Here we have presented the 2.2 Å crystal structure of the human telomeric NCP and characterised its properties in solution. Whilst the overall structure of the telomeric NCP is similar to that of other published NCP structures there are differences in details. Significantly, the overall stability of the telomeric NCP is lower than that of other NCPs and it is more dynamic than canonical nucleosomes. These differences likely impact on the biological role of telomeric chromatin. Besides this, the Telo-NCP structure presented is the only high-resolution NCP structure containing a natural DNA sequence that through its repetitive DNA sequence lacks nucleosome positioning information.

Human telomeric DNA consists of tandem arrays of the six bp repeat TTAGGG. The 6 bp sequence periodicity is out of phase with the required rotational phasing close to that of the DNA helicity (∼10 bp) (22). The present high-resolution crystal structure of the Telo-NCP containing a145 bp telomeric DNA achieves its sequence positioning on the HO by symmetric stretching on both sides of the dyad. This is made possible by several flexible TA and TT base steps located in inward-facing minor grooves as well as in adjacent locations in inward-facing regions. The structural adjustments of these steps enable the bending necessary for wrapping around the HO permitting the important interactions between DNA phosphates and the surface of the H3/H4 tetramer. The out-of-phase periodicity of the telomeric DNA sequence also results in the positioning of rigid GG and GT steps at inward-facing minor grooves which is accommodated by a pronounced deformation of these base steps. Since such distortion requires significant energy, the necessary positioning of the GG/GT steps might account for the low stability, affinity and increased DNA breathing that we have observed for the Telo-NCP. The observation that telomeric DNA containing NCPs easily form well-defined, highly-diffracting crystal reaffirms the observation that most DNA sequences form nucleosomes (78,79). Indeed, the very first crystals of nucleosomes were obtained from NCPs isolated from natural sources having a complex mixture of sequences (2). Hence, the present work opens up the perspective to crystalise and determine the high-resolution structure of NCPs of other biologically relevant weak or non-positioning DNA sequences. The telomeric nucleosome is also an ideal starting point for studying shelterin components like TRF1 and TRF2 and their interactions with the telomeric nucleosome.

Telomeres are hotspots for DNA damage, and the elevated instability and dynamics of the telemetric nucleosome that we observed in solution, combined with the fact that the telomeric TTAGG repeat is particularly prone to damage (80), helps to rationalise the DNA damage accumulation at telomeres. Furthermore, DNA damage response (DDR) remodels chromatin near the damage location and telomeres are highly susceptible to DDR (8,9), Our observations of lower stability, as well as similar stretching in both Telo-NCP structures, suggest that telomeric nucleosomes are ideal substrates for chromatin remodelers, which rebuild chromatin in one base pair sub-steps (81) and this could explain their high susceptibility to remodelling. Additionally, telomeres in vivo are characterised by dynamic epigenetic states displaying both euchromatic and heterochromatic marks that show variation between cell lines (82). Together with the previous observations of dynamic nucleosomes in telomeric chromatin (23,24), our observations suggests that that the dynamic properties of the telomeric nucleosome imparted by its unique DNA sequence, helps mediate rapid disassembly of compact chromatin states, which facilitates recruitment of chromatin-modifying enzymes that introduce epigenetic marks thereby contributing to the dynamic regulation of epigenetic states at telomeres.

Our data reveal the high-resolution structure and dynamic solution properties, which has important implications for the biological properties of the telomeric nucleosome. These findings have implications for our understanding of the dynamics of the human telomere, and therefore for our insight into normal and aberrant telomere function. With this new knowledge, we can now progress to answer longstanding questions in telomere biology, including the mechanism of telomeric nucleosome packing into compact chromatin structures and of the interactions with telomere-specific DNA binding factors with telomeric chromatin.

## DATA AVAILABILITY

Atomic coordinates have been deposited in the Protein Data Bank under accession codes PDB 6KE9 (Telo-NCP single orientation), PDB 6L9H (Telo-NCP-1, orientation-1) and PDB 6LE9 (Telo-NCP-2, orientation-2). Other data are available from the authors upon request.

## SUPPLEMENTARY DATA

Supplementary Data are available at NAR online.

## ACKNOWLEDGEMENTS

We are grateful to the Swiss Light Source (SLS), Villigen PSI, Switzerland for providing X-ray synchrotron beamtime at the X06DA (PXIII) beamline, and The National Synchrotron Radiation Research Center (NSRRC) at Hsinchu, Taiwan is acknowledged for allocation of beamtime that enabled the Synchrotron X-ray scattering measurements. We are indebted to Dr Chun-Jen Su (NSRRC) for technical assistance. We are highly indebted to Curt Davey for comments and suggestions throughout this project and to Simon Lattman and Wu Bin for valuable discussions.

## FUNDING

This research was supported by a Singapore Ministry of Education Academic Research Fund (AcRF) Tier 3 (MOE2012-T3-1-001) and Tier 2 (MOE2018-T2-1-112) grants. We also acknowledge the NTU Institute of Structural Biology (NISB) for supporting this research.

## Author contributions

AS, CWL, HLT, NK, DR and LN designed the experiments. AS, CWL, HLT, NB and VO carried out the experiments. AS, CWL, NB and NK analyzed the data. All authors reviewed drafts of the manuscript and AS, DR and LN wrote the paper.

## COMPETING INTERESTS

The authors declare no competing interests.

## SUPPLEMENTARY DATA

**Supplementary Figure S1.**
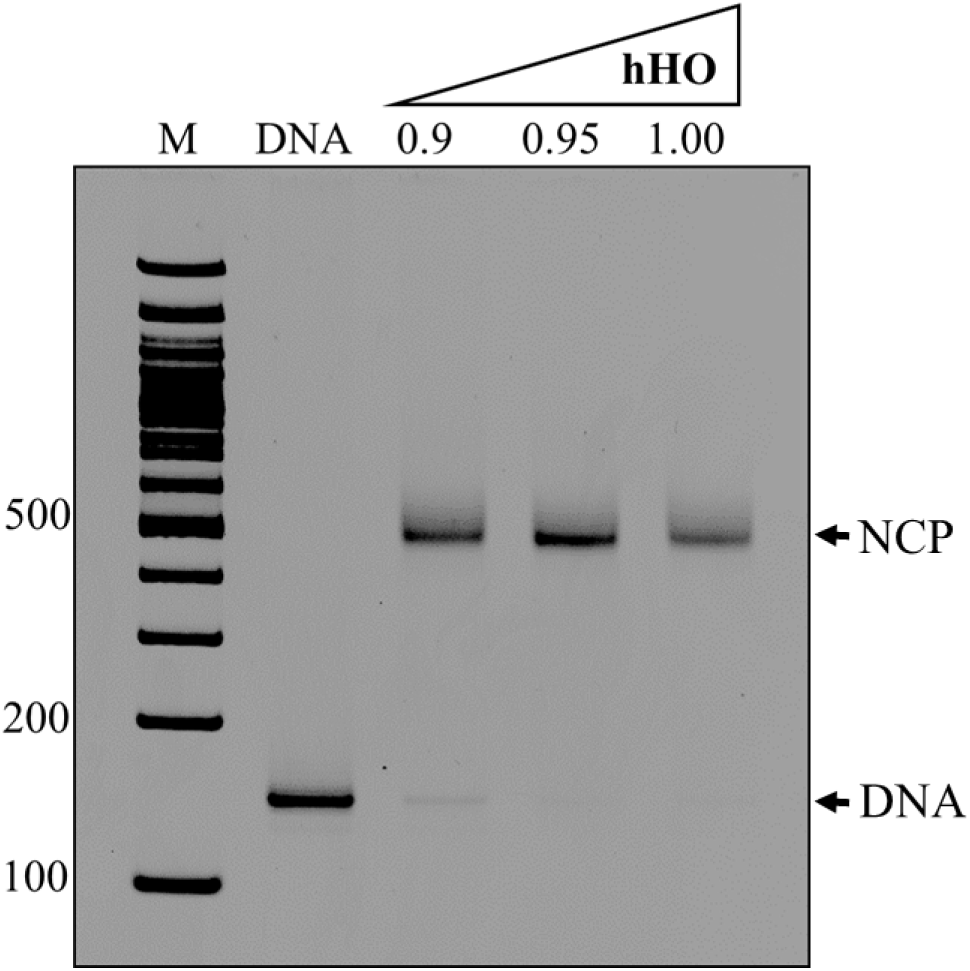
Reconstitution of Telo-NCP. Lane M is DNA marker, lane DNA is reference 145 bp telomeric DNA and lanes 0.9, 0.95 and 1.00 show products of the Telo-NCP reconstitution with the respective human HO:DNA ratios.

**Supplementary Table S1.**
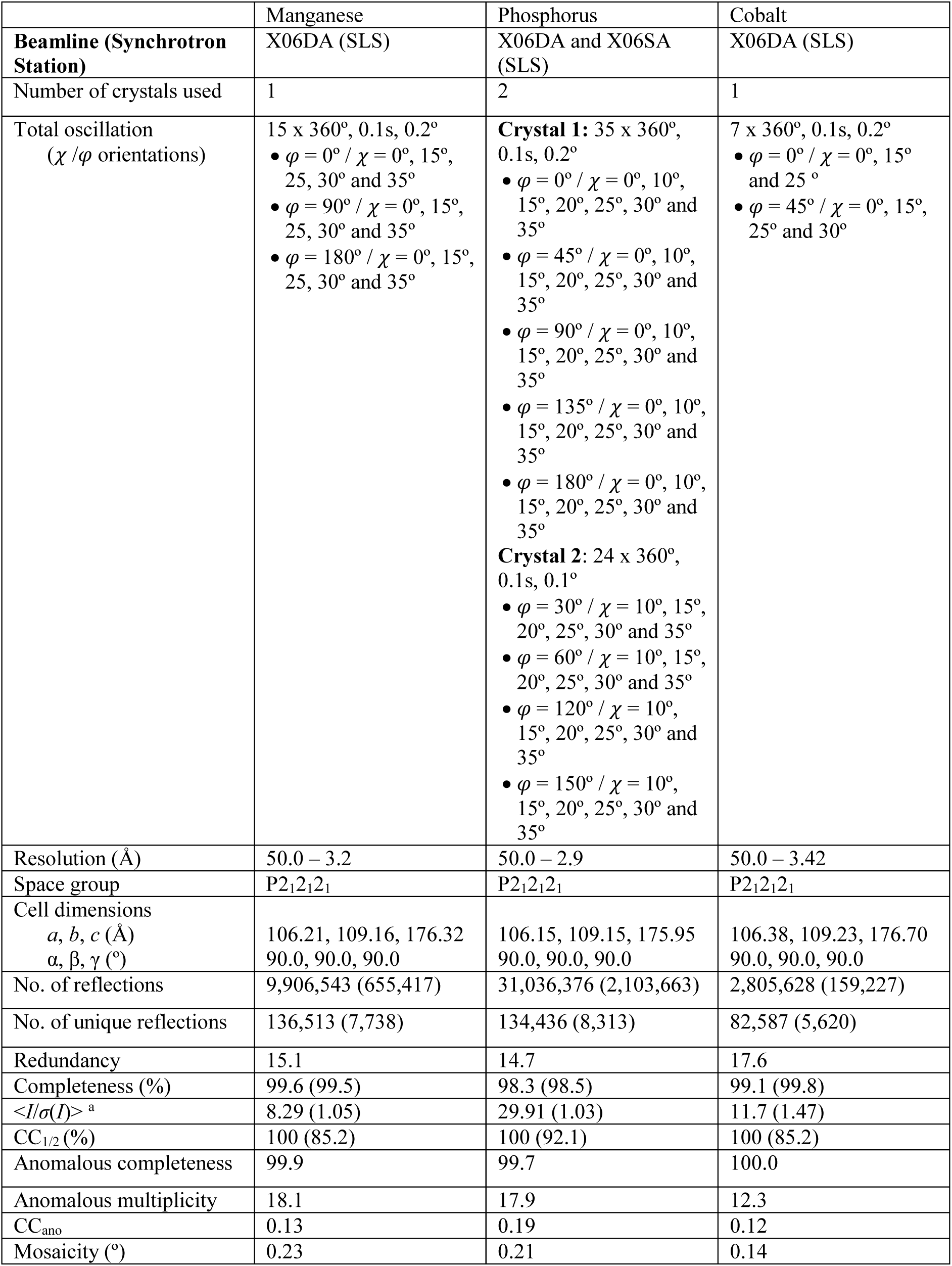
Anomalous data statistics for the telomeric NCP (dataset 1, two orientations).

**Supplementary Figure S2.**
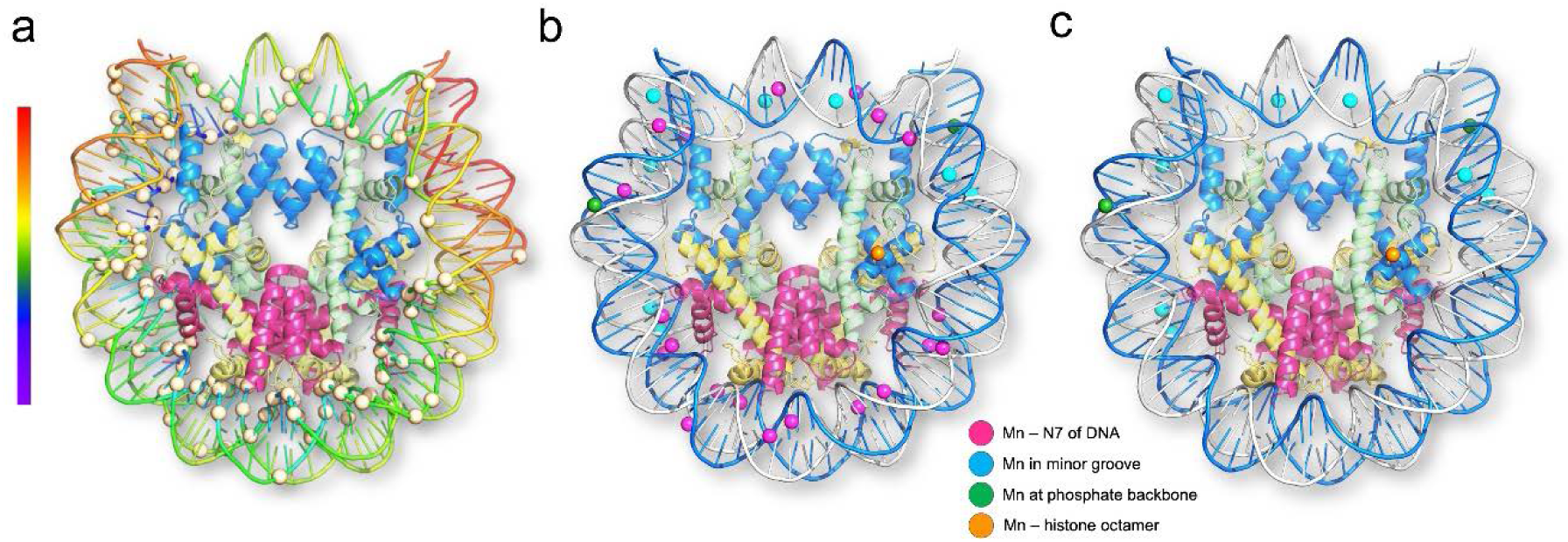
A solution of the structure of the two-orientation telomeric NCP. The initial structure was obtained by molecular replacement and by rebuilding the bases from DNA phosphate location as inferred from the phosphorous anomalous map (**a**) Location of phosphorous atoms determined from the phosphorous anomalous map. 114 phosphorous atoms (light yellow spheres) that can be confidently located were used to build the DNA structure. The DNA is plotted by temperature factors. Refinement of the rebuilt structure resulted in diffused densities for the bases of the DNA indicating averaging over two orientations. To verify this, we collected manganese and cobalt single-wavelength anomalous datasets (Supplementary Table S1). Mn^2+^ and Co^2+^ ions coordinate with N7 and O6 of guanine in ‘GG’ or ‘GC’ steps of the major groove and in the telomeric nucleosomes will only bind to one of the DNA strands. **(b)** The overall distribution of Mn^2+^ ions on the telomeric NCP. Mn^2+^ coordination with the N7 of guanine is shown as pink spheres. (**c**) Distribution of Mn^2+^ ions that do not coordinate with the N7 of guanine. Mn^2+^ ions located near the phosphate backbone, histone octamer and minor groove are displayed as green, orange and blue spheres respectively.

**Supplementary Figure S3.**
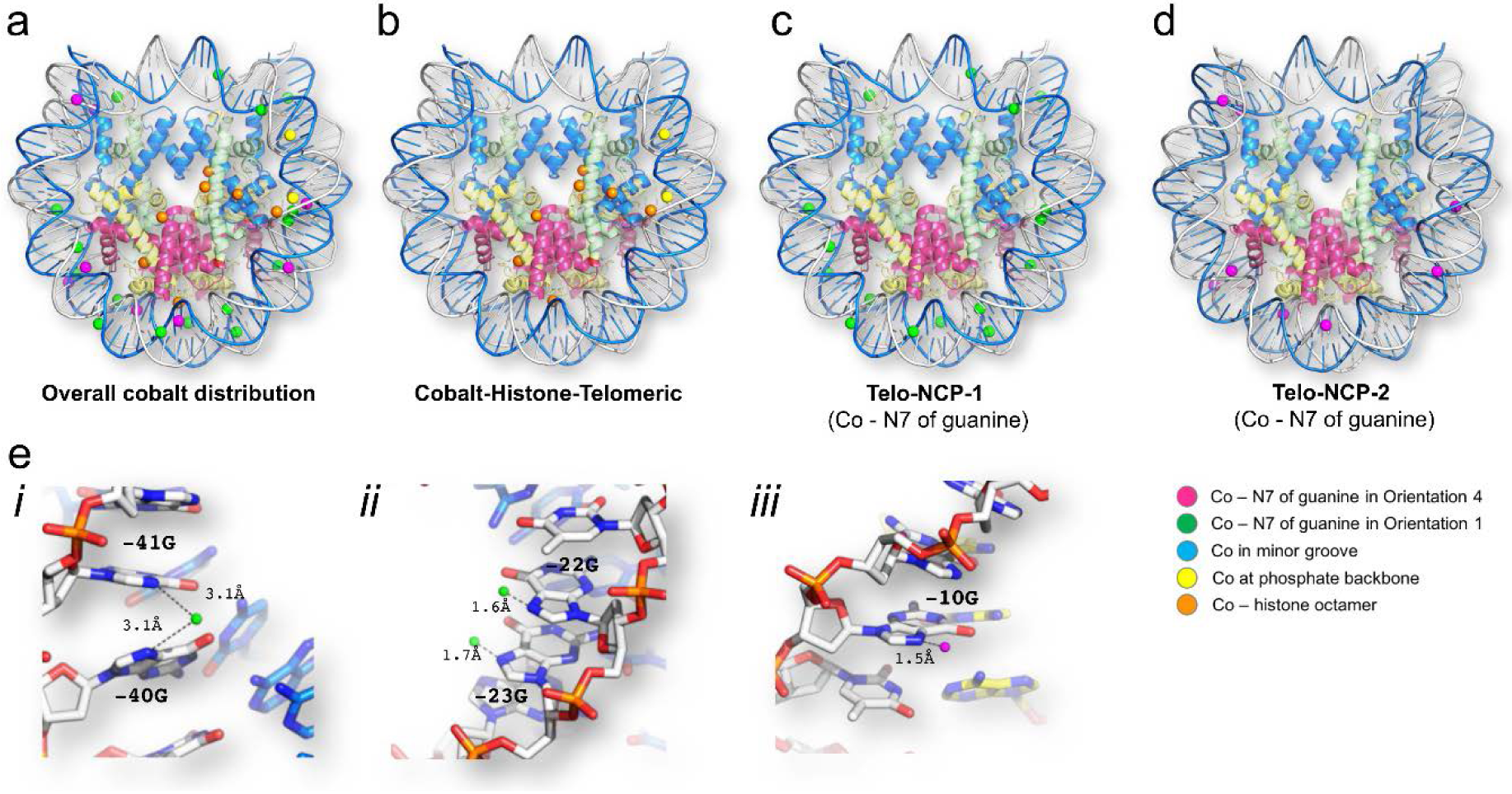
Locations of the Co^2+^ ions in the Telo-NCP. **(a)** The overall distribution of Co^2+^in the Telo-NCP. Co^2+^ coordination to the guanines N7 in the major groove of the Telo-NCP-1 is shown as magenta spheres and those interacting with the guanine N7 in the Telo-NCP-2 are depicted as green spheres. Co^2+^ coordination to the phosphate backbone, histone octamer and minor groove is shown as yellow, orange and blue spheres respectively. **(b)** Co^2+^coordination to the histone octamer and phosphate backbone. **(c)** Co^2+^ coordination to the guanine N7 in the Telo-NCP-1. **(d)** Co^2+^ interaction with guanine bases in the Telo-NCP-2. **(e)** Two modes of Mn^2+^ coordination to the guanine N7 are also observed for Co^2+^. Examples of Co^2+^ coordination with the single guanine N7 and with the two adjacent guanine N7 atoms are shown.

**Supplementary Table S2.**
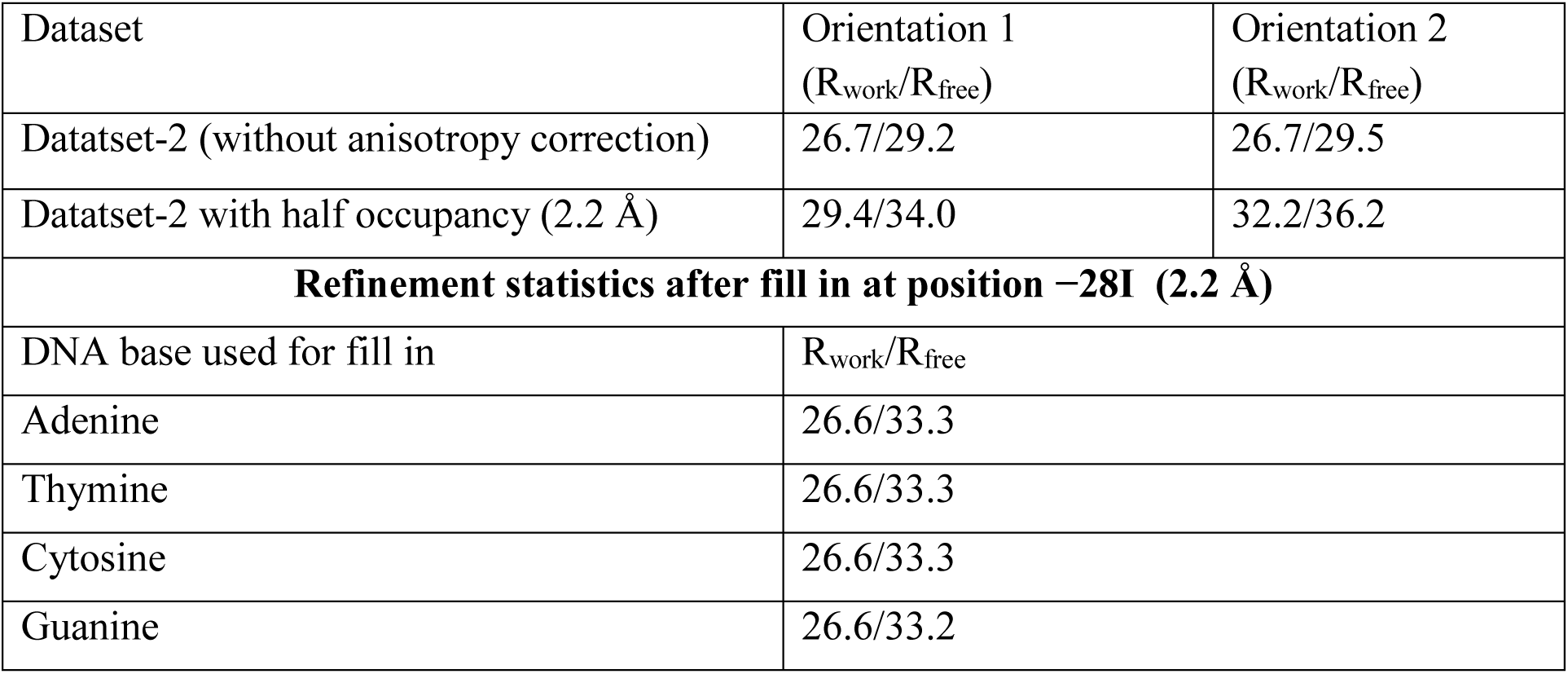
Refinement statistics of the dataset-2 with a single orientation of the NCPs in the crystal lattice. Refinements were carried out to investigate the preference for single orientation with full occupancy (without anisotropy correction) and half occupancy (with anisotropy correction). Without anisotropy correction, we did not notice significant preference, but with half occupancy and anisotropy correction, there was a clear preference for orientation 1. Omit maps were also created with deletion of guanine bases, followed by refinement with filling in with the four DNA bases. The refinement statistics and the maps from one such deletion of base −28I is shown below and in Supplementary Figure S4 below.

| Dataset | Orientation 1<br>($R_{\text{work}}/R_{\text{free}}$) | Orientation 2<br>($R_{\text{work}}/R_{\text{free}}$) |
| --- | --- | --- |
| Datatset-2 (without anisotropy correction) | 26.7/29.2 | 26.7/29.5 |
| Datatset-2 with half occupancy (2.2 Å) | 29.4/34.0 | 32.2/36.2 |
| **Refinement statistics after fill in at position –28I (2.2 Å)** |  |  |
| DNA base used for fill in | $R_{\text{work}}/R_{\text{free}}$ | |
| Adenine | 26.6/33.3 |  |
| Thymine | 26.6/33.3 |  |
| Cytosine | 26.6/33.3 |  |
| Guanine | 26.6/33.2 |  |

**Supplementary Figure S4.**
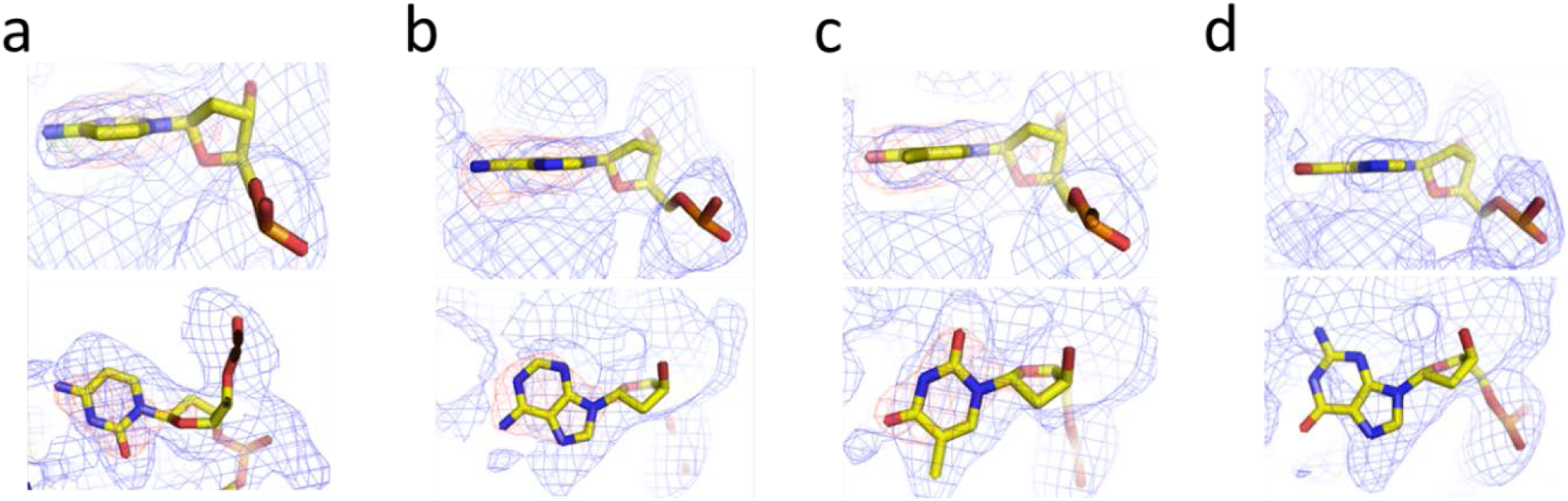
Refinement with omit maps indicating the presence of the NCP in single orientation in the dataset 2. Omit maps were generated by deletion of guanine bases (an example for the −28I position is shown). The four DNA bases cytosine (a) adenine (b), thymine (c) and guanine (d) were used to ‘fill in’ for the missing nucleotide and refined. The side view and top view of the nucleotide at position −28I after the refinements superimposed with the blue (named 2Fo−Fc) (1σ) and red (Fo−Fc) (3σ) density map are shown. The maps from the refinements with adenine, thymine and cytosine display intensity in the Fo−Fc maps in (a), (b) and (c), indicative of an incorrect fit with the electron density at −28I. Refining the structure with guanine gave a correct fit, confirmed by the absence of the red density in the Fo−Fc map in (d). In an NCP crystal with two orientation, the position −28I in the second orientation will be occupied by cytosine and refinement with cytosine would not have given a Fo−Fc map density. Multiple other locations occupied by guanines were also investigated and gave similar results confirming that presence of a single orientation in dataset 2.

**Supplementary Figure S5.**
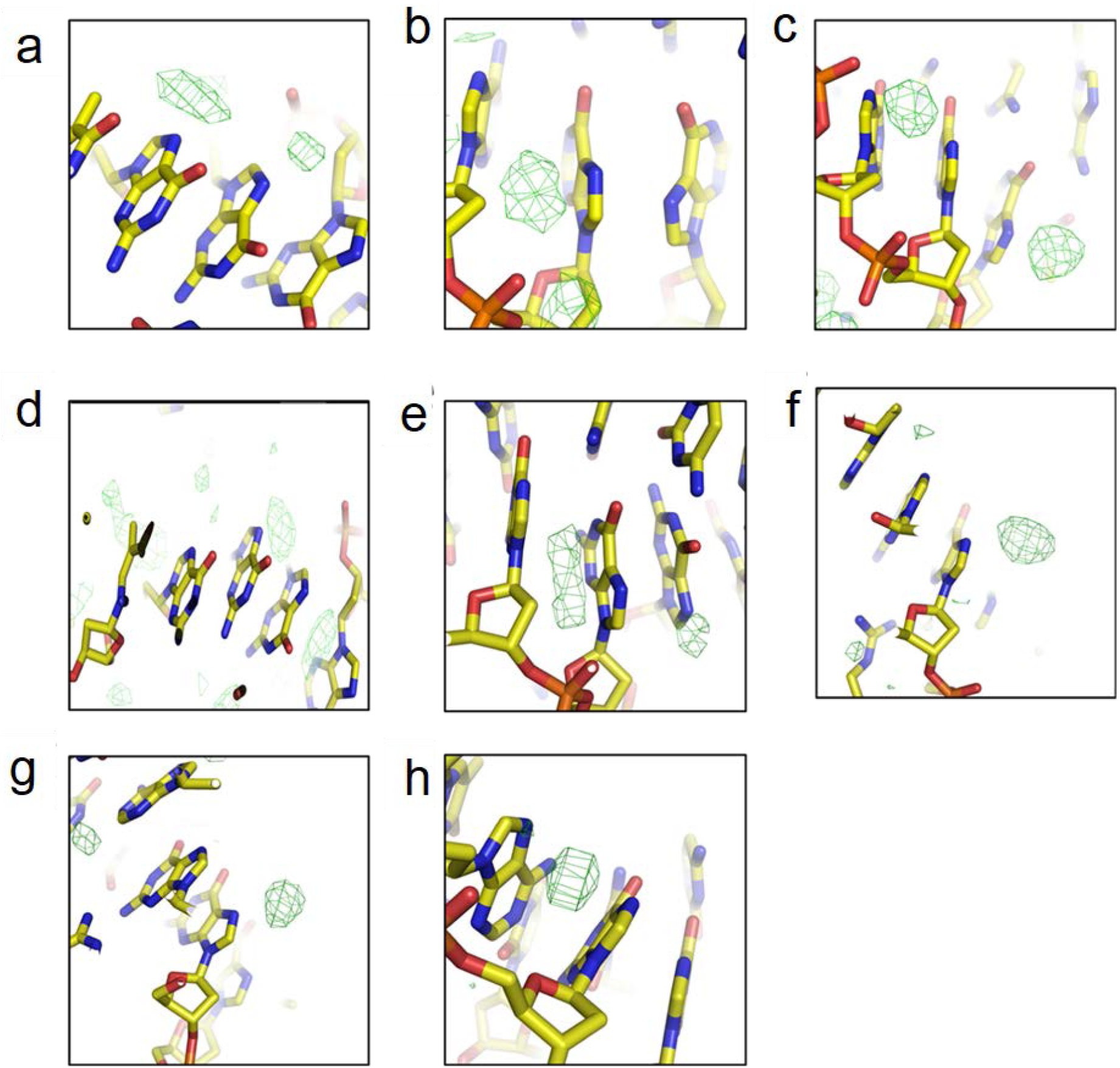
Putative Mn^2+^ electron densities (shown in green) in the Telo-NCP crystal of the dataset-2 with a suggested single NCP orientation. The putative Mn^2+^ densities are visible in the Fo−Fc map at the expected coordination distance near the guanine N7. These signals at coordination distance from N7 of guanine were investigated for both orientations. The densities are stronger (higher σ) and more numerous in the dataset-2 comparing to the similar signals observed in the electron density map of the dataset-1 where two orientation of the Telo-NCP is observed (see Fig. S6). This is a confirmation that most of the NCPs are in a single orientation in the dataset 2. The locations of putative signals in the Telo-NCP orientation 1, dataset 2 are presented at the positions **(a)** −64I, −65I and −66I. **(b)** −13I, −12I, −11I. **(c)** −5I, −6I, −7I. **(d)** 5I, 6I, 7I. **(e)** 24I, 25I 26I. **(f)** 35I, 36I, 37I. **(g)** 48I, 49I 50I. **(h)** 60I, 61I and 62I. The image is rendered with coot (Fo−Fc map −3σ)

**Supplementary Figure S6.**
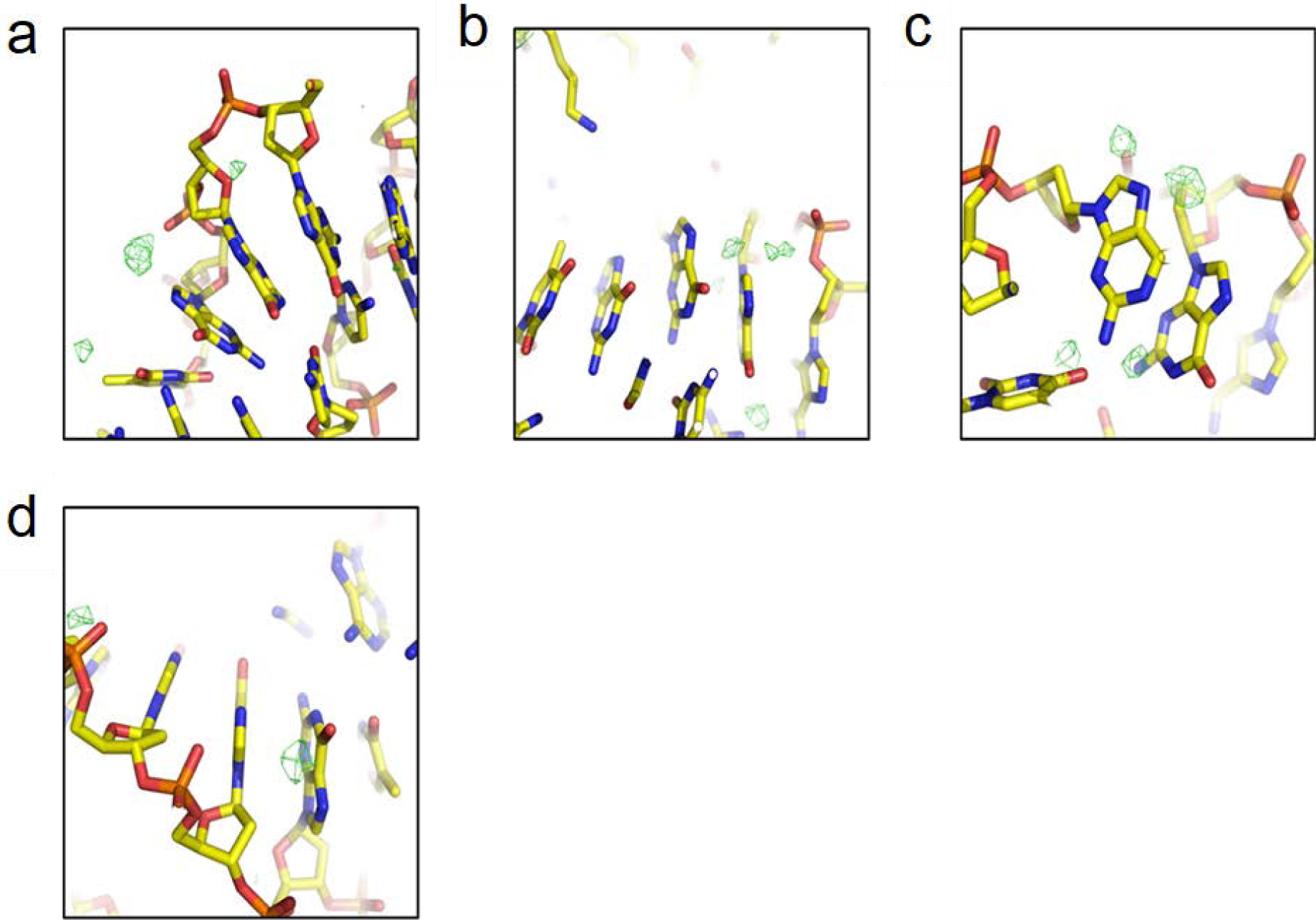
Putative Mn^2+^ electron densities for the Telo-NCP in orientation 2 of the dataset-2. The signals at coordination distance from N7 of guanine of the orientation 2 were fewer in number and weaker. The locations of putative signals (green blobs) in orientation 2 are presented **(a)** −28J. **(b)** 7J. **(c)** 13J. **(d)** 38J. (Fo−Fc map −3σ).

**Supplementary Figure S7.**
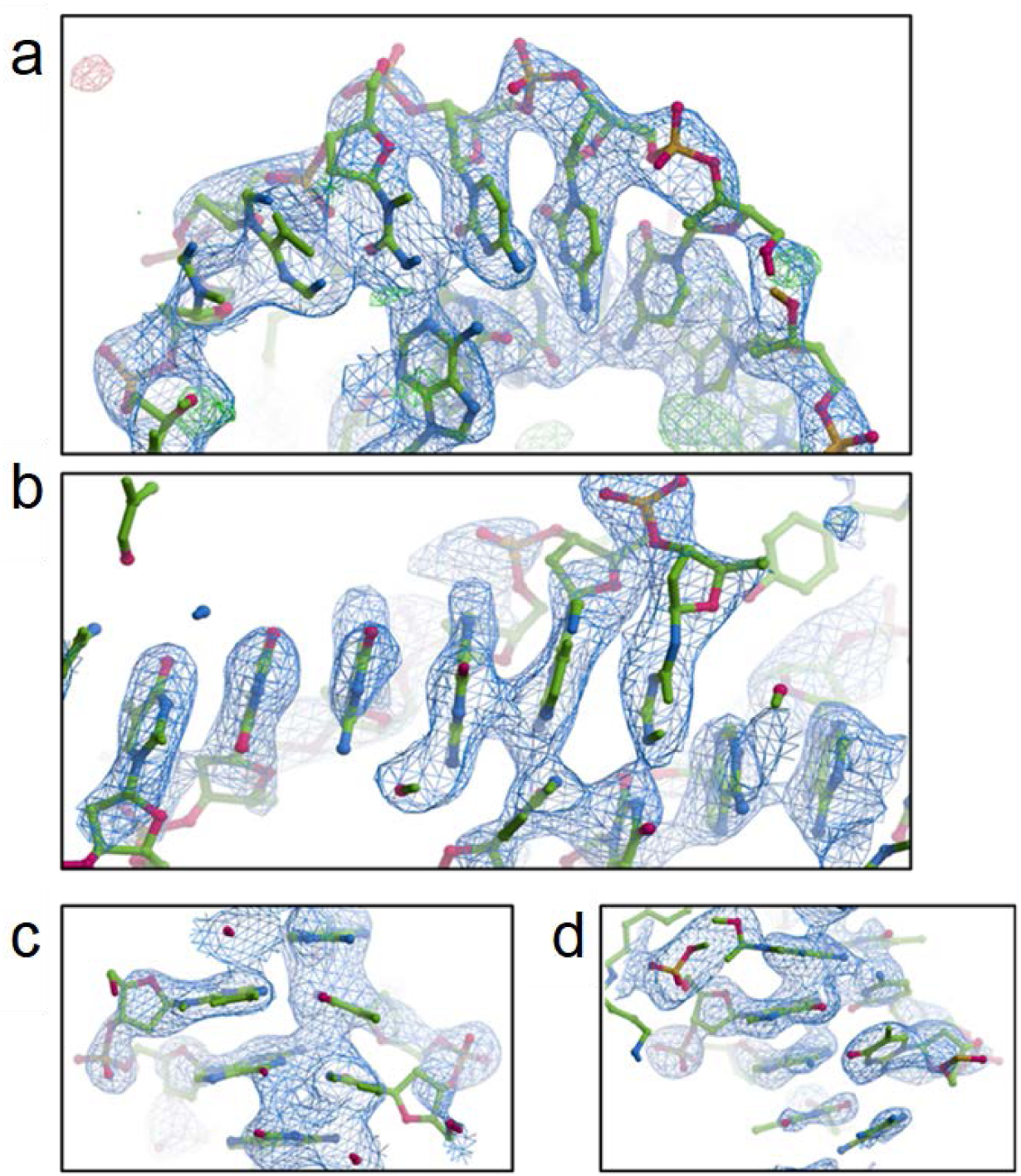
Snapshots from the electron density map of 2.2 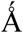 Telo-NCP structure. The DNA from dataset-2 shows well-defined electron density with base separation. Map rendered with COOT (1) at a contour level of 1.49σ. (**a**) SHL 2. (**b**) SHL 6.5. (**c**) Dyad. (**d**) SHL −7. Note: The map shown is from refinements carried out with REFMAC.

**Supplementary Figure S8.**
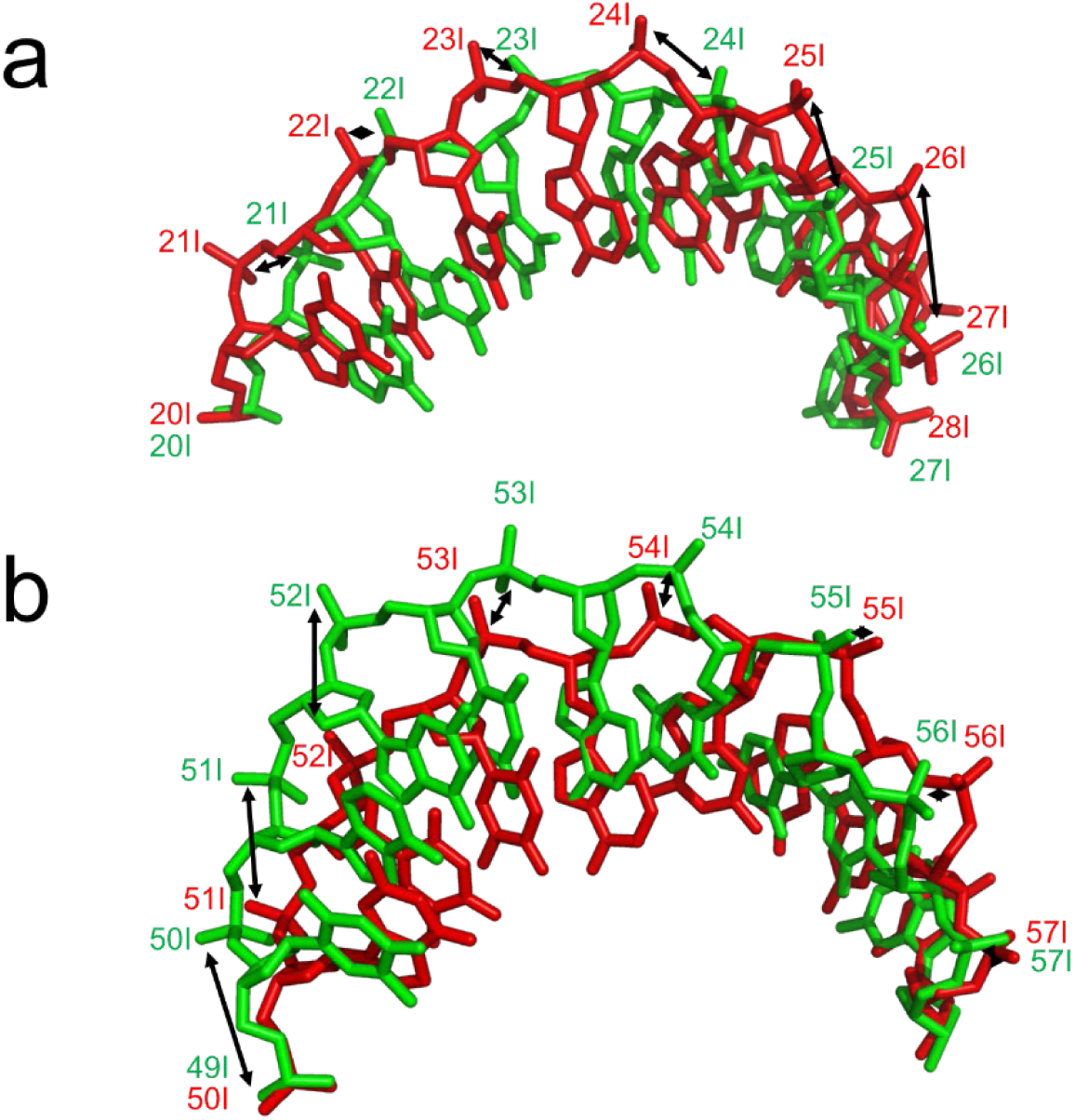
Comparison of DNA path in the Telo-NCP crystal with the palindromic 145 bp alpha satellite sequence (α-sat-NCP, PDB code 2NZD (1)). Telo-NCP is shown in red, the α-sat-NCP structure is in green. There are two regions of deviation, the deviations in the positive half of NCP is shown (SHL 0-7). **(a)** The stretching of DNA in the α-sat-NCP from 20I to 27I (SHL 2 to SHL 2.5) is missing in Telo-NCP. The double-headed black arrow indicates the deviation between corresponding phosphates. The DNA phosphates of the two NCPs are in phase until 20I. The stretching across bases 20I to 28I is characterized by increasing difference in positions of the corresponding bases. By the base pair 28I, the phosphates are out of phase by one bp with respect to the corresponding atoms of the α-Sat-NCP. The DNA remains out of phase by one base pair from 28I to bp 50I (SHL 2.5 to SHL 5). **(b)** The stretch across 50I to 57I (SHL to SHL 5.5) by one bp on the Telo-NCP brings the phosphate back in phase with the phosphates of the alpha satellite NCP.

**Supplementary Table S3.** Base steps at pressure points where the minor groove faces inward for the Telo-NCP 2.2 Å-resolution structure as well as for the Telo-NCP-1 2.5 Å-resolution orientation 1 structure (dataset 2 with two orientations of the NCP in the crystal lattice). Comparison with α-sat NCP (1) NCP-601 (2) and NCP-601L (3). All NCPs consist of 145 bp DNA.

| SHL | Telo-NCP,<br>dataset 2, one<br>orientation | Telo-NCP,<br>dataset 1, two<br>orientations | α-sat NCP<br>(2NZD) | 601 NCP<br>(3LZ0) | 601L NCP<br>(3UT9) |
| --- | --- | --- | --- | --- | --- |
| -5.5 | TA | TA | AG | CG | CG |
| -4.5 | GG | GG | AA | TC | TC |
| -3.5 | GG | GG | TG | TA | TA |
| -2.5 | TA | TA | TC | TA | TA |
| -1.5 | TT | TT | GG | TA | TA |
| -0.5 | GG | GG | AG | TA | TA |
| 0.5 | AG | AG | CT | CC | TA |
| 1.5 | TT | TT | CC | TA | TA |
| 2.5 | GG | GG | GA | GG | TA |
| 3.5 | AG | AG | CA | CC | TA |
| 4.5 | TA | TA | TT | AG | GA |
| 5.5 | GT | GT | CT | TC | CG |

**Supplementary Figure S9.**
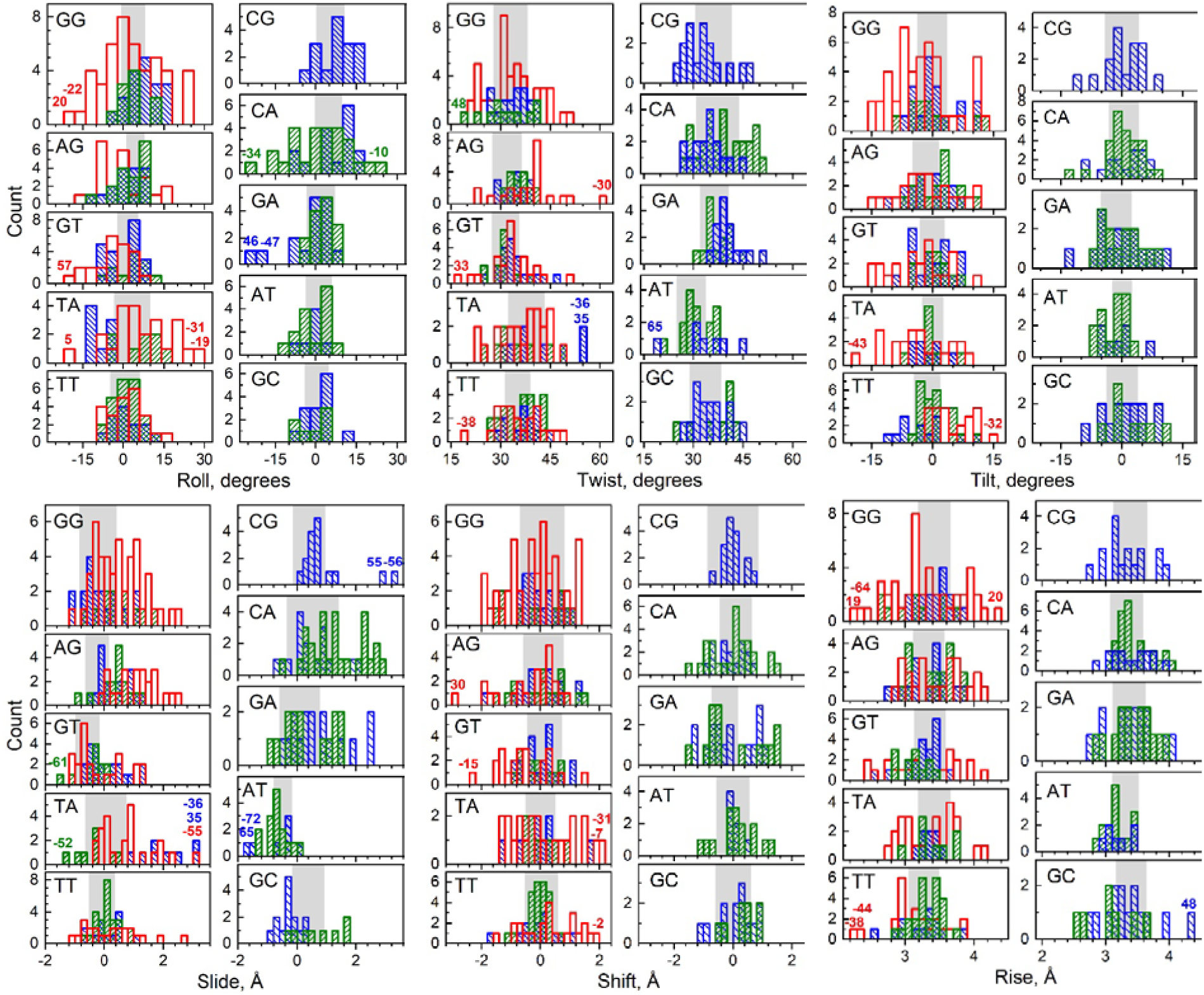
Population distribution of base-pair step parameters in the Telo-NCP in comparison with the same steps present in the NCP-601L and α-satellite NCP. Base steps from Telo-NCP, NCP-601L and α-satellite NCP are shown respectively as red, blue and green columns. In each panel, the shaded area covers the range of values (mean ± s.d.) observed in DNA crystals according to the analysis by Olson et al (4). Coloured numbers mark base-pair step positions of extreme deviation of the respective parameters. It may be noted that the Telo-NCP GT base step 15 exhibits an extreme tilt value of 36 degrees and is outside the display range and not shown in the corresponding figure. Some of the more pronounced deformation cases observed in the Telo-NCP are illustrated in Supplementary Figure S10 below.

**Supplementary Figure S10.**
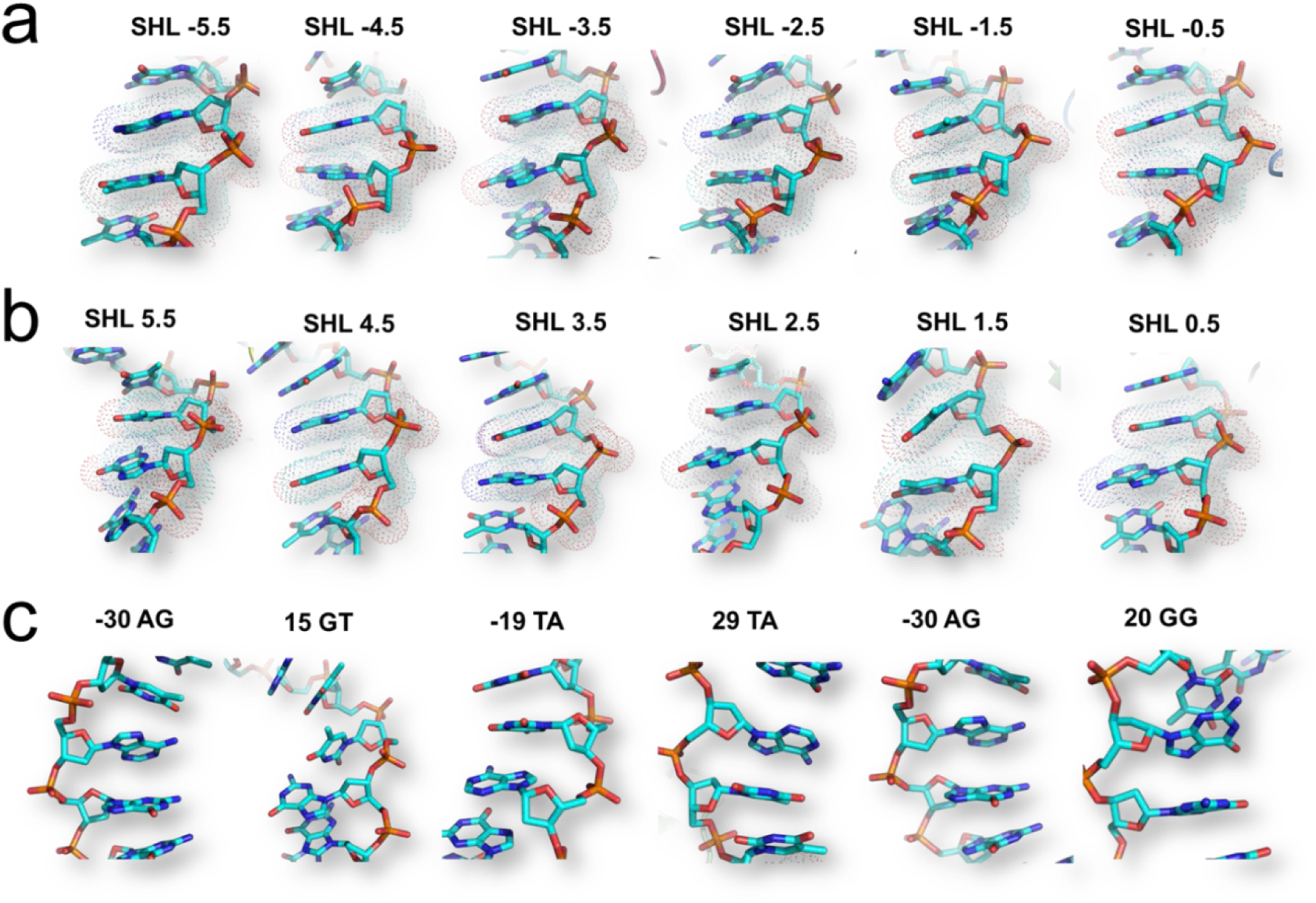
Base step deviations in Telo-NCP. The DNA is shown as sticks and the base steps at the pressure points in the minor groove are highlighted with dots. (**a**) The base steps at pressure points in the positive half of the NCP SHL 0 to 5.5. (**b**) The base steps at pressure points in the negative half of the NCP SHL 0 to −5.5. (**c**) Examples of structures of base-pair steps that show highly pronounced deviation from the parameters of an ideal B-form DNA.

**Supplementary Figure S11.**
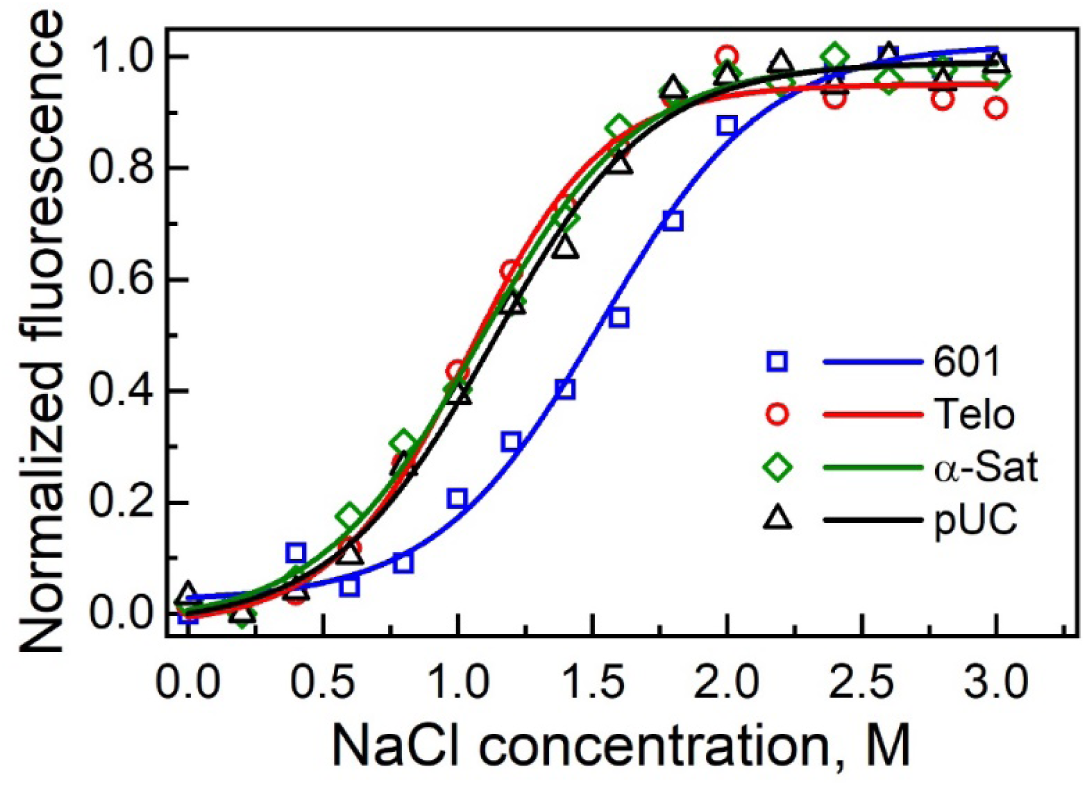
Salt-dependent stability of different NCPs in NaCl buffer (20 mM Tris (pH 7.5), 1 mM EDTA, 1 mM DTT, NaCl was varied from 0 to 3 M) measured by tyrosine fluorescence at 305 nm. The 601-NCP is shown in blue, Telo-NCP in red, Telo-NCP bound with cisplatin in yellow, α-sat-NCP in green and pUC-NCP in black. Each curve represents an average of three runs. Five measurements were obtained for each data point in individual runs. In reconstitutions of all NCPs, the same batch of histone octamer was used.

**Supplementary Figure S12.**
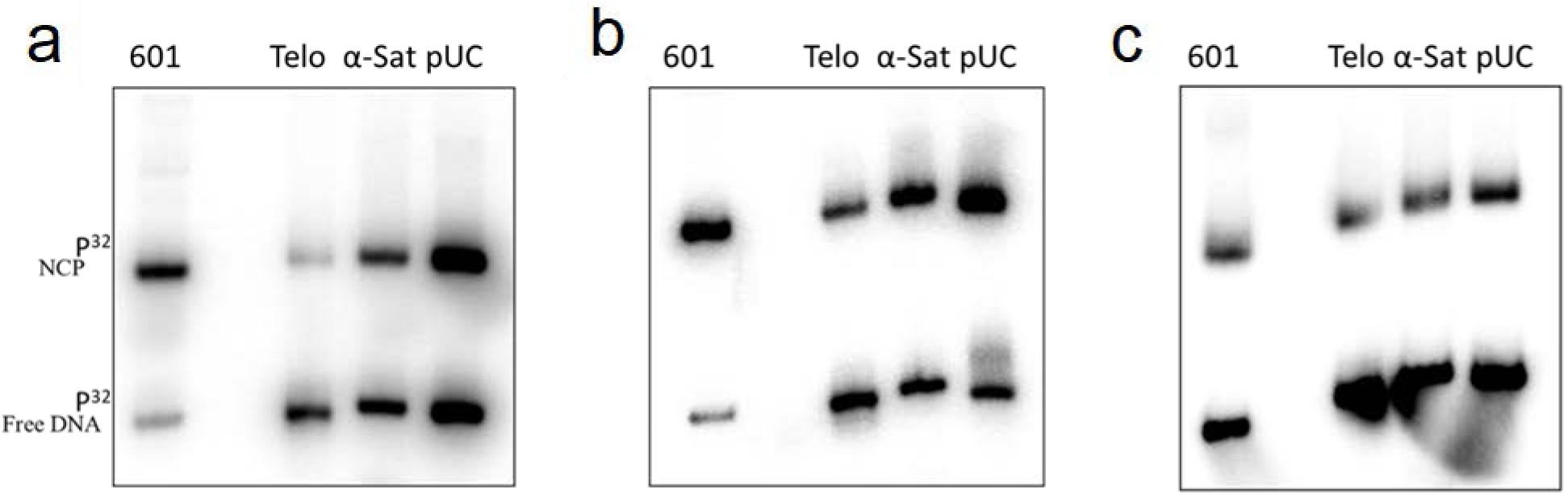
Competitive NCP reconstitution. In all gel images, lanes marked 601, Telo, α-sat and pUC show respective reconstituted NCPs and radiolabelled DNA not included in the NCP. **(a)** The PAGE analysis of competitive reconstitution in LiCl using radiolabelled DNA. In salt dialysis, most of the labelled 601 DNA is reconstituted into NCP. In comparison with 601 DNA, pUC and α-sat lanes show a higher ratio of free DNA to NCP indicating that a lower proportion of DNA was incorporated into the NCP. The ratio of free DNA to NCP is lowest for telo DNA suggesting that the least amount of telomeric DNA incorporated into NCP. **(b)** The PAGE analysis of competitive reconstitution in LiCl supplemented with 1 mM MgCl_2_ using radiolabelled DNA. With the addition of Mg^2+,^ there is a minor increase in the amount of DNA incorporated into NCP for all four DNA sequences. **(c)** The PAGE analysis of competitive reconstitution carried out using radiolabelled DNA employing Nap1 and ACF after 4 hours incubation. In contrast to salt dialysis, only a small fraction of the DNA is incorporated into nucleosome for all four DNA.

**Supplementary Figure S13.**
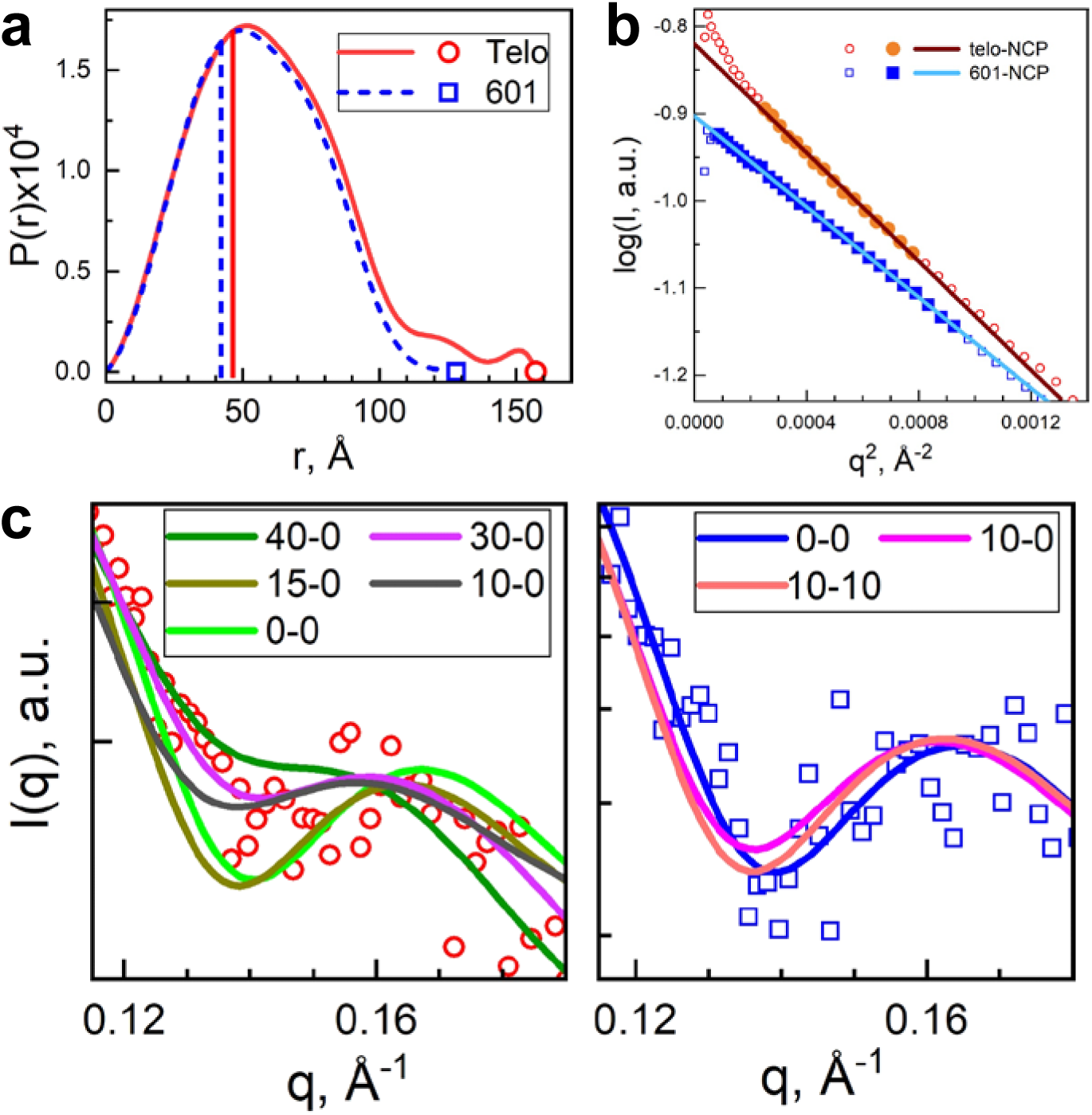
Solution SAXS studies of Telo-NCP and 601-NCP. **(a)**. Comparison of the distance distribution functions, P(r), calculated from the solution SAXS profiles of Telo-NCP (1 mg/mL, red symbol and lines) 601-NCP (1 mg/mL, blue symbol and dashed lines). Vertical lines indicate the values of the corresponding R_g_. The points indicate the values of D_max_ (maximal pair distance) of each NCP type. Numerical values are given in Supplementary Table S4. (**b**). Example of Guinier plots calculated from the SAXS spectra of the Telo-NCP and 601-NCP solutions shown in Fig. 7 of the main text. Points used for linear fitting are displayed as solid symbols; calculated Rg values are given in Supplementary Table S4. (**c**). Comparison of SAXS profiles recorded at 1 mg/mL NCP concentration at low salt (10 mM KCl) for the Telo-NCP (left panel) and 601-NCP (right panel). Experimental data are shown as points and form factors calculated from molecular structures as curves. For the modelled curves, two numbers show the lengths of DNA base pairs unwrapped form the entry/exit of the NCP. For the Telo-NCP the form factors of the NCPs with asymmetric DNA positioning or unwinding of up to 40 bp give the best fitting to the experimental data. For the 601-NCP, the best fit is observed with the NCP structures showing no or small distortion from the fully wrapped DNA conformation. Numerical data is given in Supplementary Table S4.

**Supplementary Table S4.** Comparison of the outputs from SAXS profiles obtained for the Telo-NCP and 601-NCP in experiment and simulated from modelled molecular structures, including the radius of gyration *(R*_*g*_) and the maximal intra-atom distances within a particle (*D*_*max*_) determined from the distance distribution function *P(r)*. The top row of numbers shows parameters obtained from the experiment; two sections below present output of the analysis of molecular structures constructed from stretches of straight DNA and NCP structure generated from crystal structures or from MD simulations. Best fittings of the modelled form factors to the experimental SAXS profiles are highlighted by bold font.

| Number of unwrapped bp | $R_g$ from Guinier plot, Å | $R_g$ from $P(r)$, Å | $D_{max}$, Å | $\chi^2$, fit quality, |
| --- | --- | --- | --- | --- |
| Data extracted from SAXS measurement 601-NCP, 1.0 mg/mL |  |  |  |  |
| Experiment | **42.1** +/- 0.1 | **42.1** +/- 0.1 | **128** | 0.986, excellent |
| NCP crystal structures |  |  |  |  |
| 3LZ0 | 39.8 | 39.9 | 117 | 8.15 |
| Telo-NCP | 39.7 | 38.8 | 116 | 8.56 |
| NCP structures with collapsed tails generated in MD simulations |  |  |  |  |
| **0-0** | 41.6 | 40.8 | 117 | **1.05** |
| **10-0** | 42.5 | 42.1 | 132 | **1.69** |
| **10-10** | 42.4 | 42.0 | 132 | **1.74** |
| 15-0 | 42.9 | 42.5 | 131 | 2.09 |
| 20-0 | 44.0 | 43.6 | 138 | 2.87 |
| 15-15 | 44.1 | 43.7 | 143 | 3.50 |
| Data extracted from SAXS measurement Telo-NCP, 1.0 mg/mL |  |  |  |  |
| Experiment | **46.3** +/- 0.2 | **46.4** +/- 0.2 | **157** | 0.840, good |
| NCP crystal structures |  |  |  |  |
| 3LZ0 | 39.8 | 39.9 | 117 | 15.26 |
| Telo-NCP | 39.7 | 38.8 | 116 | 16.72 |
| NCP structure with collapsed tails generated in MD simulations |  |  |  |  |
| **40-0** | 50.4 | 50.3 | 199 | **1.44** |
| **30-0** | 46.1 | 45.8 | 166 | **1.66** |
| 0-0 | 41.1 | 40.8 | 130 | 2.57 |
| 10-0 | 41.9 | 41.6 | 132 | 2.66 |
| 15-0 | 42.0 | 41.8 | 131 | 2.66 |
| 25-0 | 44.4 | 44.1 | 154 | 2.65 |
| 10-10 | 42.1 | 41.8 | 132 | 3.20 |
| 15-15 | 43.4 | 43.1 | 143 | 3.81 |
| 20-20 | 43.6 | 43.4 | 140 | 4.36 |

## Notes

#### Summary of Updates

Revisions in response to reviewers comment during revision process at journal peer review stage.

